# Biomarker discovery in progressive supranuclear palsy from human cerebrospinal fluid using mass spectrometry-based proteomics

**DOI:** 10.1101/2023.10.05.561075

**Authors:** Yura Jang, Sungtaek Oh, Anna J. Hall, Zhen Zhang, Thomas F. Tropea, Alice Chen-Plotkin, Liana S. Rosenthal, Ted M. Dawson, Chan Hyun Na, Alexander Y. Pantelyat

**Author notes:** These authors made equal contributions (Yura Jang, Sungtaek Oh).

## Abstract

Progressive supranuclear palsy (PSP) is a neurodegenerative disorder that is often misdiagnosed as Parkinson’s Disease (PD) because of shared symptoms. PSP is characterized by the accumulation of tau protein in specific brain regions, which results in loss of balance, gaze impairment, and dementia. Diagnosing PSP is often challenging, and there’s a significant demand for reliable biomarkers. However, existing biomarkers, including tau protein and neurofilament light chain (NfL) levels in cerebrospinal fluid (CSF), show inconsistencies in distinguishing PSP from other neurodegenerative disorders. To overcome these limitations, we conducted a comprehensive proteome analysis for CSF samples from 40 PSP, 40 PD, and healthy controls (HC) using the tandem mass tag-based quantification method, identifying 3,653 unique proteins. Our statistical analysis identified 190, 152, and 247 differentially expressed proteins when comparing PSP vs. HC, PSP vs. PD, and PSP against both PD and HC, respectively. Gene set enrichment analysis and interactome analysis conducted with the differentially expressed proteins in PSP CSF indicated that most of them were implicated in cell adhesion, cholesterol metabolism, and glycan biosynthesis. Cell-type enrichment analysis revealed that neuronally-derived proteins were predominant among the differentially expressed proteins. Potential biomarker classification performance showed that ATP6AP2 (reduced in PSP) had the highest AUC (0.922), followed by NEFM, EFEMP2, LAMP2, CHST12, FAT2, B4GALT1, LCAT, CBLN3, FSTL5, ATP6AP1, and GGH. This is the first large-scale mass spectrometry-based proteome analysis to discover CSF PSP biomarkers differentiating from both controls and PD, thereby laying a foundation for further development and validation.

## INTRODUCTION

Progressive supranuclear palsy (PSP) is a neurodegenerative Parkinsonian disorder with an estimated prevalence of about 5 to 6 in every 100,000 people worldwide, typically beginning after the age of 60.^1–3^ The pathological features of PSP are characterized by progressive accumulation of 4-repeat tau, formation of globose neurofibrillary tangles, and neuronal loss in the brainstem, basal ganglia, and cortex.^1^ Two of the classic clinical signs of patients with PSP are impairment of vertical gaze and balance loss with backward falls.^4^ The most common initial symptom of people with PSP is balance loss^5^ and subsequently, PSP patients show changes in their mood and behavior and develop dementia over time.^1–3^ It is recognized that PSP can have different clinical presentations, and the current clinical diagnostic criteria include several variants in addition to the most common PSP-Richardson syndrome.^6^

The clinical evaluation of early-stage and variant forms of PSP is challenging, with limited sensitivity and specificity, making it difficult to distinguish PSP from alternative diagnoses such as Parkinson’s Disease (PD).^6^ In recent years, diagnostic approaches have evolved, exploiting magnetic resonance imaging (MRI) and positron emission tomography (PET).^7^ Currently, there are no established diagnostic biomarkers of PSP, owing to observed discrepancies between clinical manifestations and underlying neuropathological findings. These inconsistencies hinder their utilization in key areas such as early-stage diagnosis, precise pathological characterization, and longitudinal tracking of disease progression.^8^ Further research and development are essential for discovering and optimizing PSP biomarkers, given their potential importance in understanding and managing the disease.

Multiple research groups have endeavored to establish an accurate diagnosis of PSP by identifying specific CSF biomarkers.^1,4^ Since the abnormal accumulation of tau proteins within brain cells is considered a potential target for developing therapeutic interventions for PSP, the predominant studies on CSF biomarkers for PSP were focused on tau proteins.^9–11^ However, the relationship between total tau, phosphorylated tau and tau fraction levels in CSF and the disease’s clinical presentation may be complex, and not necessarily distinct from healthy control (HC) groups in certain contexts.^2,12^

On the other hand, multiple studies report that neurofilament light chain (NfL) concentrations are 2 to 5 times higher in the CSF of PSP patients compared to HC and Parkinson’s Disease (PD) groups, and similar results were observed in plasma.^3–5^ Nevertheless, the diagnostic specificity of NfL for PSP remains inconclusive.^3,4,13,14^ To tackle this challenge, in this study, we conducted a mass spectrometry-based proteomics experiment for the identification of additional biomarkers in CSF of PSP patients. We analyzed 120 CSF samples from 40 PSP, 40 PD and 40 control individuals. We exploited 11-plex tandem mass tags (TMTs) to analyze 120 samples more accurately. This study represents a comprehensive mass spectrometry-based proteomic analysis of human CSF from PSP patients, aiming to identify PSP biomarkers that distinguish it from PD and controls. The candidate biomarkers discovered in this study—if validated—will pave the way for the development of reliable PSP biomarkers.

## METHODS

### Collection of cerebrospinal fluid samples

We employed CSF samples from 40 PSP, 40 PD, and 40 healthy older control (HC) individuals well-matched on gender and age. The CSF samples were collected from study volunteers at the University of Pennsylvania using the previously described Parkinson’s Disease Biomarkers Program CSF collection protocol^15^ (procedure Manual: https://biosend.org/docs/studies/PDBP/PDBP%20Manual%20of%20Procedures.pdf). Briefly, the CSF samples were collected from study participants in polypropylene vials, spun down at 2000 x g for 10 minutes at room temperature (18°C to 25°C), aliquoted, and stored at ‒80°C. Samples were shipped on dry ice to Johns Hopkins and stored at ‒80°C. The sample information is provided in Table 1. This study was approved by the University of Pennsylvania Institutional Review Board. Informed consent was obtained from each participant at study enrollment in accordance with the Declaration of Helsinki.

**Table 1.** Demographic information of the CSF samples used in this study.

|  | Diagnosis |  |  | Total (N=120) | Test |
| --- | --- | --- | --- | --- | --- |
|  | HC (N=40) | PD (N=40) | PSP (N=40) |  |  |
| Age.Yrs |  |  |  |  | P value: 0.0253<br>(Kruskal-Wallis<br>rank sum test) |
| Mean (95% CI) | 67.7 (65.5; 69.9) | 64.1 (61.7; 66.5) | 68.8 (66.6; 71.0) | 66.9 (64.5; 69.2) |  |
| Median | 66 | 65 | 69 | 66.5 |  |
| Gender |  |  |  |  | P value: 0.1864<br>(Pearson's Chi-<br>squared test) |
| Female | 27 (67.5%) | 19 (47.5%) | 24 (60%) | 70 (58.3%) |  |
| Male | 13 (32.5%) | 21 (52.5%) | 16 (40%) | 50 (41.7%) |  |
| Race |  |  |  |  | P value: 4.3e-05<br>(Pearson's Chi-<br>squared test) |
| Black | 10 (25%) | 1 (2.5%) | 0 (0%) | 11 (9.2%) |  |
| White | 26 (65%) | 39 (97.5%) | 39 (97.5%) | 104 (86.7%) |  |
| More than one<br>race | 4 (10%) | 0 (0%) | 0 (0%) | 4 (3.3%) |  |
| American<br>Indian | 0 (0%) | 0 (0%) | 1 (2.5%) | 1 (0.8%) |  |
| Education.Yrs |  |  |  |  | P value: 0.0038<br>(Kruskal-Wallis<br>rank sum test) |
| Mean (95% CI) | 15.1 (16.5; 13.8) | 17.1 (17.7; 16.5) | 14.9 (15.8; 14) | 15.7 (16.7; 14.7) |  |
| Median | 16 | 18 | 14.5 | 16 |  |

CI: confidence interval

### Sample preparation for the mass spectrometry analysis

We conducted mass spectrometry analysis of 120 CSF samples using 13 batches of 11-plex tandem mass tag (TMT, Thermo Scientific) experiments. For the normalization of data from 13 batches of the TMT experiment, we included the master pool (MP) in the last channel of each batch. We also included quality control (QC) in 10 batches to monitor data quality. To minimize the batch effect, the batch allocation and the order of 120 CSF samples and QCs were block-randomized, keeping diagnosis, sex, and age balanced using an in-house R-script. The MP was created by mixing equal volumes from all 120 CSF samples. The CSF used for QC came from a control CSF, separate from the 40 other control CSFs. The MP was divided into each batch after completing the TMT labeling. The QC was divided into 10 batches before reduction and alkylation. Two hundred twenty microliters of each CSF sample were used in this study. All CSF samples, including QC and MP, were prepared by adding 1 volume of 10 M urea in 100 mM triethylammonium bicarbonate (TEAB; Sigma). To perform the reduction and alkylation, 10 mM tris (2-carboxyethyl) phosphine hydrochloride (TCEP; Thermo Scientific) and 40 mM chloroacetamide (CAA; Sigma) were added in the CSF samples and then incubated at room temperature for 1 h. Protein digestion was carried out using LysC (Lysyl endopeptidase mass spectrometry grade; Fujifilm Wako Pure Chemical Industries Co., Ltd., Osaka, Japan) at the ratio of 1:50 for 3 h at 37°C and then using trypsin (sequencing grade modified trypsin; Promega, Fitchburg, WI, USA) at the ratio of 1:50 at 37°C overnight (for 15 h to 18 h) after diluting the concentration of urea from 5 M to 2 M by adding 50 mM TEAB. Peptides were purified using C_18_ Stage-Tips (3M Empore^TM^;3M, St. Paul, MN, USA) after acidifying them with trifluoroacetic acid (TFA; Thermo Scientific). The eluted solution containing peptides was vacuum-dried with a Savant SPD121P SpeedVac concentrator (Thermo Scientific). The digested peptides were labeled with 11-plex TMT reagents following the manufacturer’s instructions (Thermo Scientific). MP was labeled by TMT channel 131C, and the rest of the peptide samples were labeled by one of TMT channels 126, 127N, 127C, 128N, 128C, 129N, 129C, 130N, 130C, and 131. The labeling reaction was conducted at RT for 1 h. The remaining TMT tags were quenched by adding 100 mM tris buffer (pH 8.0; Thermo Scientific) and incubating for over 5 min at RT. The peptides for each batch were pooled and subjected to basic pH reversed-phase liquid chromatography (bRPLC) fractionation on an Agilent 1260 HPLC system (Agilent Technologies, Santa Clara, CA, USA). Briefly, the peptides were reconstituted in 10 mM TEAB and fractionated using a bRPLC column (Agilent 300 Extend-C_18_ column, 5 µm, 4.6 mm × 250 mm, Agilent Technologies) under an increasing gradient of the mobile phases consisting of 10 mM TEAB in water and 90% acetonitrile (ACN). A total of 96 fractions were collected by eluting over 97 min (the total run time: 150 min and the collection time: between 50 to 147 min) at a flow rate of 0.3 mL/min and were subsequently concatenated into 24 fractions. The eluted peptides were vacuum-dried.

### LC-MS/MS analysis

The LC-MS/MS analysis was conducted as described in previous publications with minor modifications.^16,17^ The peptide samples were analyzed on an Orbitrap Fusion Lumos Tribrid mass spectrometer interfaced with an Ultimate 3000 RSLCnano nanoflow liquid chromatography system (Thermo Scientific). The fractionated peptides were reconstituted in 0.5% formic acid (FA) and loaded onto a trap column (Acclaim™ PepMap™ 100, LC C18, 5 μm, 100 μm × 2 cm, nanoViper, Thermo Scientific) at a flow rate of 8 μL/min. Peptides were separated on an analytical column (Easy-Spray™ PepMap™ RSLC C18, 2 μm, 75 μm × 50 cm, Thermo Scientific) at a voltage of about 2.4 kV and at a flow rate of 0.3 μL/min with mobile phases of 0.1% FA in water and in 95% ACN using a linear gradient. The total run time was 120 min. The mass spectrometer was operated in data-dependent acquisition (DDA) mode. The MS1 scan range for a survey full scan was acquired from *m/z* 300 to 1800 in the Orbitrap at a resolution of 120,000 at an *m/z* 200. The automatic gain control (AGC) target for MS1 was set as 1 × 10^6^ and the maximum injection time was set to 50 ms. The most intense ions with charge states of 2 to 5 were isolated in a 3-sec cycle, fragmented using higher-energy collisional dissociation (HCD) fragmentation with 35% normalized collision energy, and detected at a mass resolution of 50,000. The precursor isolation window was set to *m/z* 1.6 with *m/z* 0.4 of offset. The AGC target for MS/MS was set to 5 × 10^4^, and the ion filling time was set to 100 ms. The dynamic exclusion was set to 30 sec with a 7 ppm of mass tolerance. Internal calibration was carried out using the lock mass option (*m/z* 445.12002) from ambient air.

### Database searches for peptide and protein identification

Database searches were conducted as described in prior publications with minor modifications.^16,17^ The acquired MS/MS spectra were searched against a human UniProt database (released in May 2018, containing protein entries of common contaminants) using SEQUEST search algorithm in the Thermo Proteome Discoverer platform (version 2.2.0.388, Thermo Scientific). The database search parameters used were as follows. The precursor mass tolerance was set to 10 ppm and the fragment mass tolerance to 0.02 Da. The maximum missed cleavages allowed was 2. Carbamidomethyl (+57.02146 Da) at cysteine and TMT tags (+229.162932 Da) modification at the N-terminus of a peptide and lysine were set as fixed modifications. Oxidation (+15.99492 Da) of methionine was set as a variable modification. The peptides and proteins were filtered at 1% of the false discovery rate. The protein quantification was performed with the following parameters and methods. Both unique and razor peptides were used for peptide quantification, while protein groups were considered for peptide uniqueness. Reporter ion abundance was computed based on signal-to-noise (S/N) ratios, and the missing intensity values were replaced with the minimum value. The quantification value corrections for isobaric tags were disabled. The average reporter S/N threshold was set to 50. Data normalization was disabled. Protein grouping was performed with a strict parsimony principle to generate the final protein groups. All proteins sharing the same set or subset of identified peptides were grouped, while protein groups with no unique peptides were filtered out. The Proteome Discoverer iterated through all spectra and selected a peptide-spectrum match (PSM) with the highest number of unambiguous and unique peptides.

### Experimental design and statistical rationale

Experimental design and statistical analyses were performed as described previously with minor modifications.^16,17^ We conducted sample size analysis using the pwr package in R. When we wanted to detect proteins with >1.35-fold differences between groups, the required minimum sample size was 31 when the significance level was 0.0001, power was 0.8, sigma was 0.338, and delta was 0.433 (=log_2_ 1.35). The sigma value of 0.338 was derived from our in-house TMT proteomics experiments for the quantification of CSF proteins. We determined the significance level of 0.0001 based on our previous studies. When we identified several thousands of proteins, most of the proteins with *P* value < 0.0001 showed a *q*-value < 0.05. Based on this sample size analysis, we decided to use 40 samples per group. The statistical analysis of the mass spectrometry data was performed with the Perseus version 1.6.0.7 software package. The protein abundance data from 13 batches of the TMT experiments were normalized by dividing the abundance values of each protein by that of MP included in each batch. The relative abundance values for each sample were log2-transformed. We removed proteins with one or more missing values across 120 samples. To further remove batch effects, an additional normalization was conducted with the ComBat package in R. The technical variation was monitored by a coefficient of variation of QCs in each experimental batch. To access the biological variation, the signal-to-noise (S/N) ratio was calculated by dividing the standard deviation (SD) of QC by the standard deviation (SD) of the samples.

Bootstrap ROC analysis was carried out using the fbroc package in R. Sampling with replacement was repeated 500 times for the bootstrap ROC. The area under the curve (AUC) of a bootstrap ROC was computed for each sampling. Mean and SD values of AUCs from 500 ROCs were then calculated. This bootstrap ROC was repeated once again after labeling permutation. The *q*-values of bootstrap ROC-based analysis data were calculated as follows: (1) The mean AUC values for non-permuted and permuted data were sorted in descending order for proteins with mean AUCs > 0.5 and in ascending order for proteins with mean AUCs < 0.5; (2) The ratios of the protein numbers for the non-permuted data to the protein numbers for the permuted data were calculated as lowering the cutoff threshold, and the ratios were used as *q*-values.

To assess the classification performance of potential biomarkers, MetaboAnalyst software (version 5.0) was employed through both univariate and multivariate receiver operating characteristic (ROC) curve analyses. These analyses were conducted as described previously with minor modifications.^18^ For the univariate ROC analysis, a bootstrapping approach was implemented, involving 500 resampling iterations, to yield an area under the curve (AUC) mean value accompanied by a 95% confidence interval. For the multivariate ROC analysis, the partial least squares discriminant analysis (PLS-DA) classification technique, coupled with the inherent feature ranking method of PLS-DA, was used. A total of two latent variables were specified for this analysis. To initiate the multivariate analysis utilizing PLS-DA, ROC curves were generated using balanced subsampling by the Monte-Carlo cross-validation (MCCV) method. In each MCCV iteration, two-thirds of the samples were employed to appraise feature significance, while the remaining one-third served to validate the models developed in the initial phase. Subsequently, the most crucial features were used to construct biomarker classification models. This procedure was reiterated multiple times to estimate the performance metrics and confidence intervals for each respective model.

### Pathway analysis

Gene set enrichment analysis (GSEA) was performed by feeding differentially expressed proteins to the Kyoto encyclopedia of genes and genomes (KEGG) pathway analysis embedded in DAVID Knowledgebase.^19,20^

### Interactome and cell-type-specific enrichment analyses

Interactome analysis was carried out by the Search Tool for the Retrieval of Interacting Genes/Proteins (STRING) protein-protein interaction (PPI) database version 11.5 (https://string-db.org/).^21,22^ We used a full STRING network to analyze functional and physical protein associations. Cell-type enrichment analysis was conducted as described previously.^23^ P-values for the cell type enrichment were calculated using Fisher exact tests.

### Data and software availability

The mass spectrometry data from this study have been deposited to the ProteomeXchange Consortium (https://www.proteomexchange.org) via PRIDE partner repository with the dataset identifier ‘PXD041417’, project name ‘Biomarkers discovery for progressive supranuclear palsy from the human cerebrospinal fluid using mass spectrometry-based proteomics.’ Reviewers can access the dataset by using ‘’ as ID and ‘kdIwetsc’ as a password.

## RESULTS

### Quantitative proteome analysis of CSF samples

To identify differentially expressed proteins in PSP, we conducted a quantitative proteome analysis of 120 CSF samples from 40 PSP, 40 PD, and 40 HC individuals. For more accurate quantification of proteins, we exploited the TMT-based quantification method. To analyze 120 CSF samples using 11-plex TMT, we conducted 13 batches of TMT experiments. To normalize the protein abundances between the different batches, we added MP to the last channel of each batch. We also added QC to a random channel in 10 batches each to monitor quantification quality (Figure 1). We first digested CSF proteins into peptides and then labeled the resulting peptides with TMT tags as described above. For in-depth protein identification, the TMT-labeled peptides were pre-fractionated by bRPLC before mass spectrometry analysis. In total, 23,508,013 MS/MS spectra were acquired, and 2,277,905 MS/MS spectra were assigned to peptides leading to the identification of 283,975 peptides and 3,653 proteins. The number of proteins that were identified across 13 batches of the TMT experiments was 1,380, which we used for the downstream data analysis (Figure 1A). To normalize the data from 13 different batches, the intensity values of each protein were normalized by the MP samples in each batch (Figure 2B, left), and then, to remove residual batch effects, another round of normalization was conducted by the combat package in R (Figure 2B, right). To visually assess the batch effects of 13 batches of the data set before and after the combat normalization, the data were plotted on 2D PCA. Batch 2 (orange) showed the biggest batch effect before the Combat normalization, but this batch effect disappeared, and overall data showed a more evenly dispersed pattern. To further assess the quality of the data, the technical variations and S/N ratio of the normalized data were examined (Figure 2C). More than 98.7% of proteins manifested technical variations of 20% or less (Figure 2C, left). On the other hand, > 99.6% of proteins manifested S/N of 1 or higher, demonstrating the outstanding measurement precision of this TMT-based quantification experiments (Figure 2C, right).

**Figure 1.**
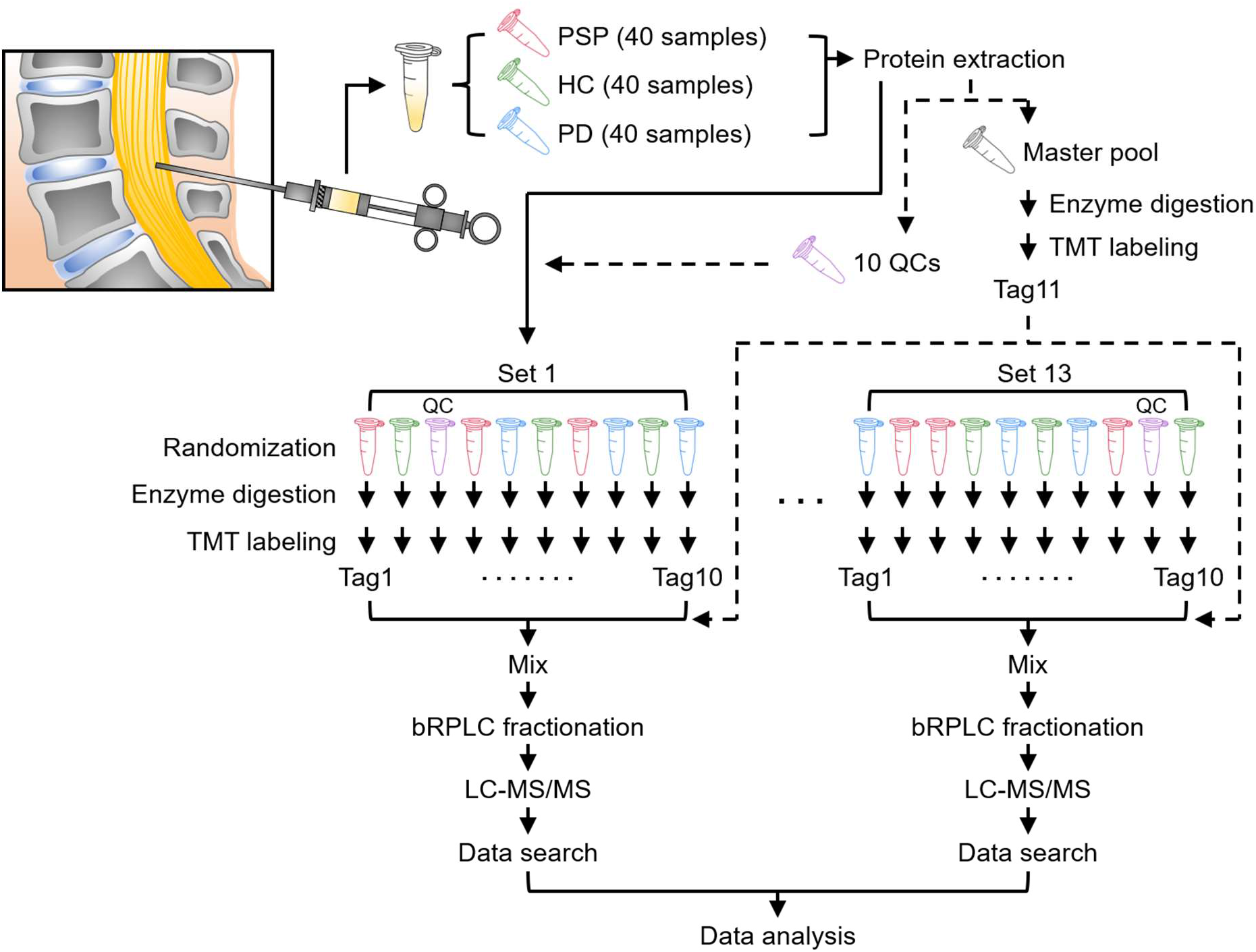
Experimental strategy for the proteomic study of the CSF samples from PSP patients, PD patients, and HC individuals. Thirteen batches of 11-plex TMT experiments were conducted to analyze the proteome of human CSF samples from 40 PSP patients, 40 PD patients, and 40 HC individuals. Master pool (MP) and QC samples were prepared by combining an equal amount of protein from all 120 CSF samples. MP was added to each batch after labeling with Tag 11 in one tube. QC was split into five aliquots and processed in each batch separately. TMT tags for individual samples and QC were determined by randomization. The proteins were digested with Lys-C and trypsin, followed by TMT labeling and prefractionation into 24 fractions prior to mass spectrometry analysis. Proteins were identified by conducting a database search of the acquired mass spectra.

**Figure 2.**
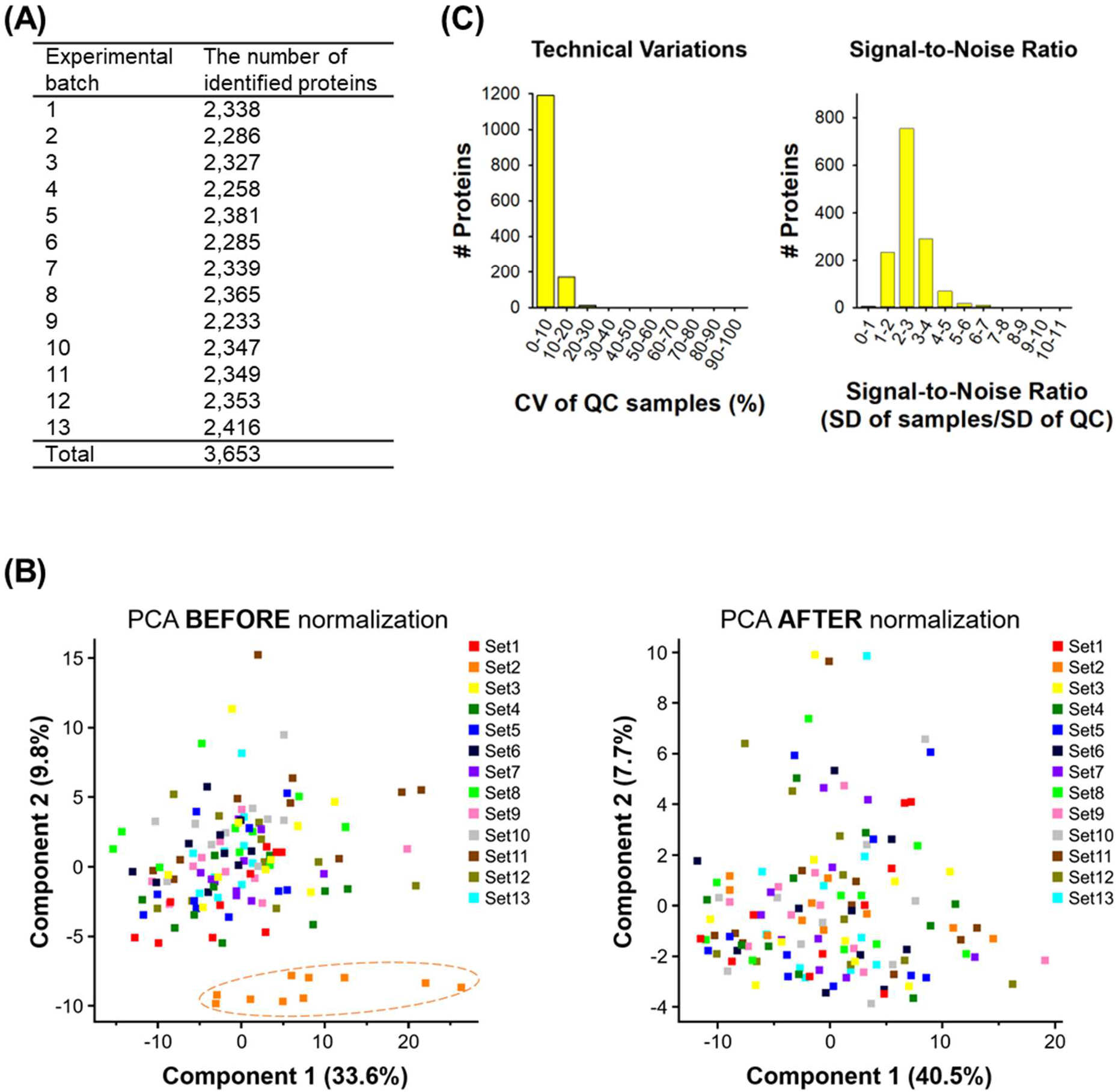
The evaluation of proteomic data quality from 13 batches of 11-plex TMT experiments. (A) The number of identified proteins in each batch. (B) To minimize batch effects of 13 different 11-plex TMT experiments, they were further normalized using the Combat package after normalizing each batch using MP. One hundred twenty CSF samples were shown on a 2D PCA plot to visualize potential batch effects before (left panel) and after (right panel) the normalization using the Combat package. (C) The coefficient of variation (CV) of QC samples and signal-to-noise ratio were calculated to evaluate data quality.

### Bootstrap ROC-based statistical analysis for the identification of differential proteins

We next conducted bootstrap ROC-based statistical analyses to identify proteins differentially expressed in PSP compared to the other groups.^16,17,24^ For the bootstrap ROC analysis, sampling with replacement was repeated 500 times generating ROC covers for each iteration. This sampling process was repeated once again after a permutation of comparison groups to estimate false discovery rate. Average AUC and SD of ROC curves were plotted (Figure 3). When we used *q*-value of < 0.01 as cutoff lines, the number of differential proteins was 190 between PSP and HC, 152 between PSP and PD, and 247 between PSP and PD plus HC. When PSP was compared to HC, NEFM was most upregulated followed by CHI3L1, SERPINA3, and MMRN1. On the other hand, ATP6AP2 showed the greatest downregulation, followed by CHST12, EFEMP2, and LAMP2 (Figure 3A). When PSP was compared to PD, a similar pattern was observed. NEFM was the most upregulated, followed by SERPINA3 and CHI3L1, while ATP6AP2 was the most downregulated, followed by EFEMP2, LAMP2, and B4GALT1 (Figure 3B). When PSP was compared to the group of PD plus HC, a similar pattern was observed. NEFM was the most upregulated, followed by SERPINA3 and CHI3L1, but ATP6AP2 was the most downregulated, followed by EFEMP2, LAMP2, CHST12, and B4GALT1 (Figure 3D). We summarized the top 50 up- and down-regulated proteins between PSP and HC (Supplemental Table S1), between PSP and PD (Supplemental Table S2), and between PSP and PD plus HC (Supplemental Table S3). Other than the top differentially expressed proteins, we also observed downregulation of NPTX2 (0.31 of mean of bootstrap AUC and 0.005 of q-value in PSP vs. PD plus HC: 0.31), which is a synaptic protein that plays a crucial role in regulating cortical network dynamics, synaptic adaptability, memory, and is associated with cognitive decline and AD progression.^25–27^ These results suggest that we successfully identified differentially expressed proteins in PSP.

**Figure 3.**
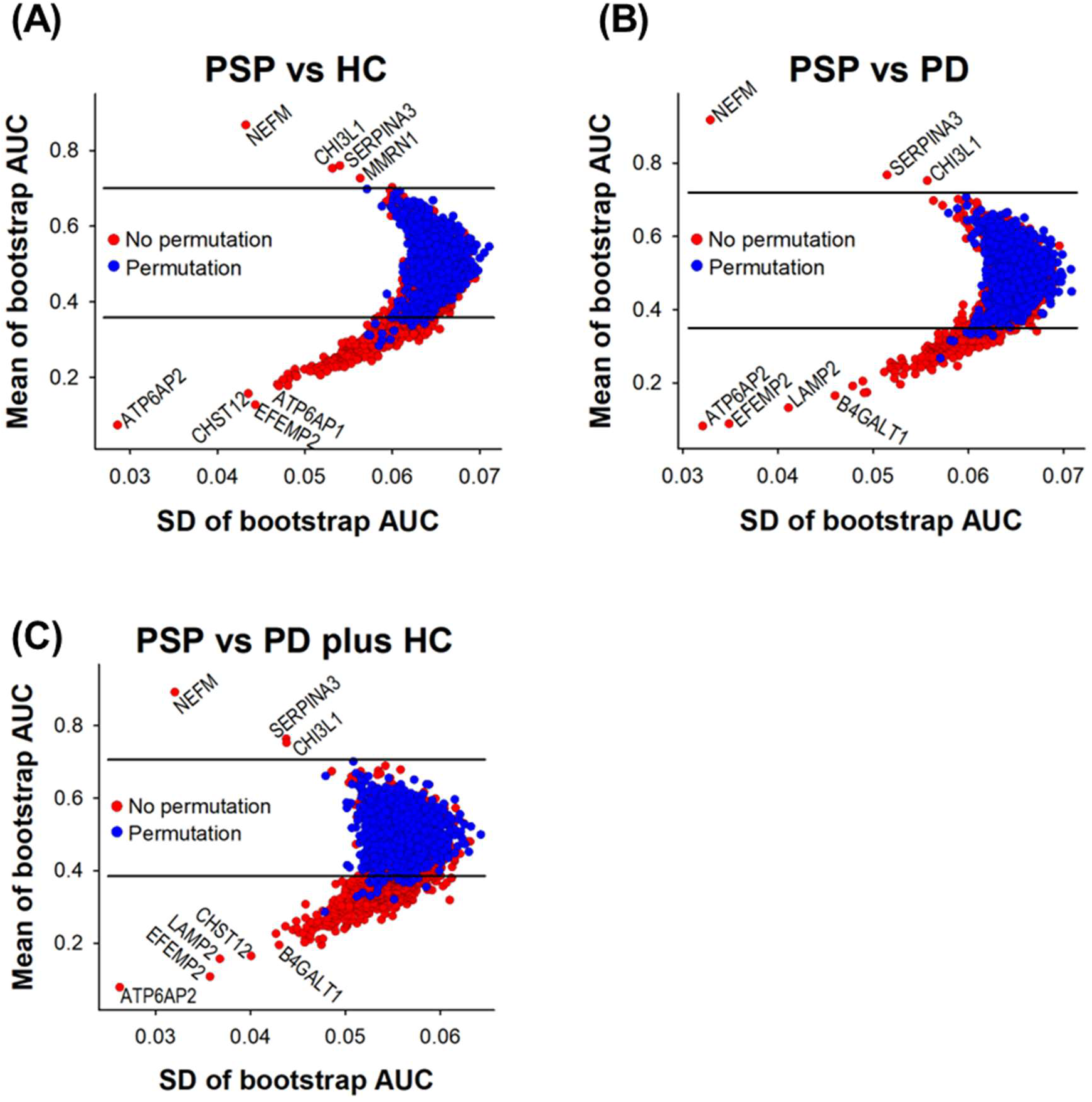
Bootstrap ROC plots of the CSF proteins identified from PSP, patients PD patients, and HC individuals. Bootstrap ROC analyses were conducted to estimate variations of resampling. To calculate *q*-values, bootstrap ROC analyses after permutation of the comparison groups were conducted too. The differentially expressed proteins with *q*-values < 0.01 are shown at the outside of the upper and lower horizontal lines. The proteins on the upper and lower side of the *q*-value line are up- and down-regulated in PSP compared to HC (A), in PSP compared to PD (B) and in PD compared to PD plus HC (C), respectively.

### Comparison of differentially expressed proteins in CSF with those from globus pallidus

The main goal of this study was to discover potential PSP biomarkers. Therefore, we needed to narrow down the list of the differentially expressed proteins in CSF. If the differentially expressed proteins in CSF reflect the changes in the brain, this change should be observed in the brain. Thus, we compared the list of differentially expressed proteins in CSF with the differentially expressed proteins in globus pallidus (GP) of PSP patients, which we reported previously.^17^ When PSP was compared to HC, 4 differentially expressed proteins overlapped between CSF and GP (Figure 4A). When PSP was compared to PD, only 2 proteins overlapped (Figure 4B). When PSP was compared to PD plus HC, 8 proteins were overlapping (Figure 4C). CNTNAP2 and EPDR1 were common differentiating proteins for PSP vs. HC and PSP vs. PD plus HC. HAPLN4 was the common differentiating protein for PSP vs. PD and PSP vs. PD plus HC. GGH was the common differentiating protein in all three comparisons (Table 2).

**Figure 4.**
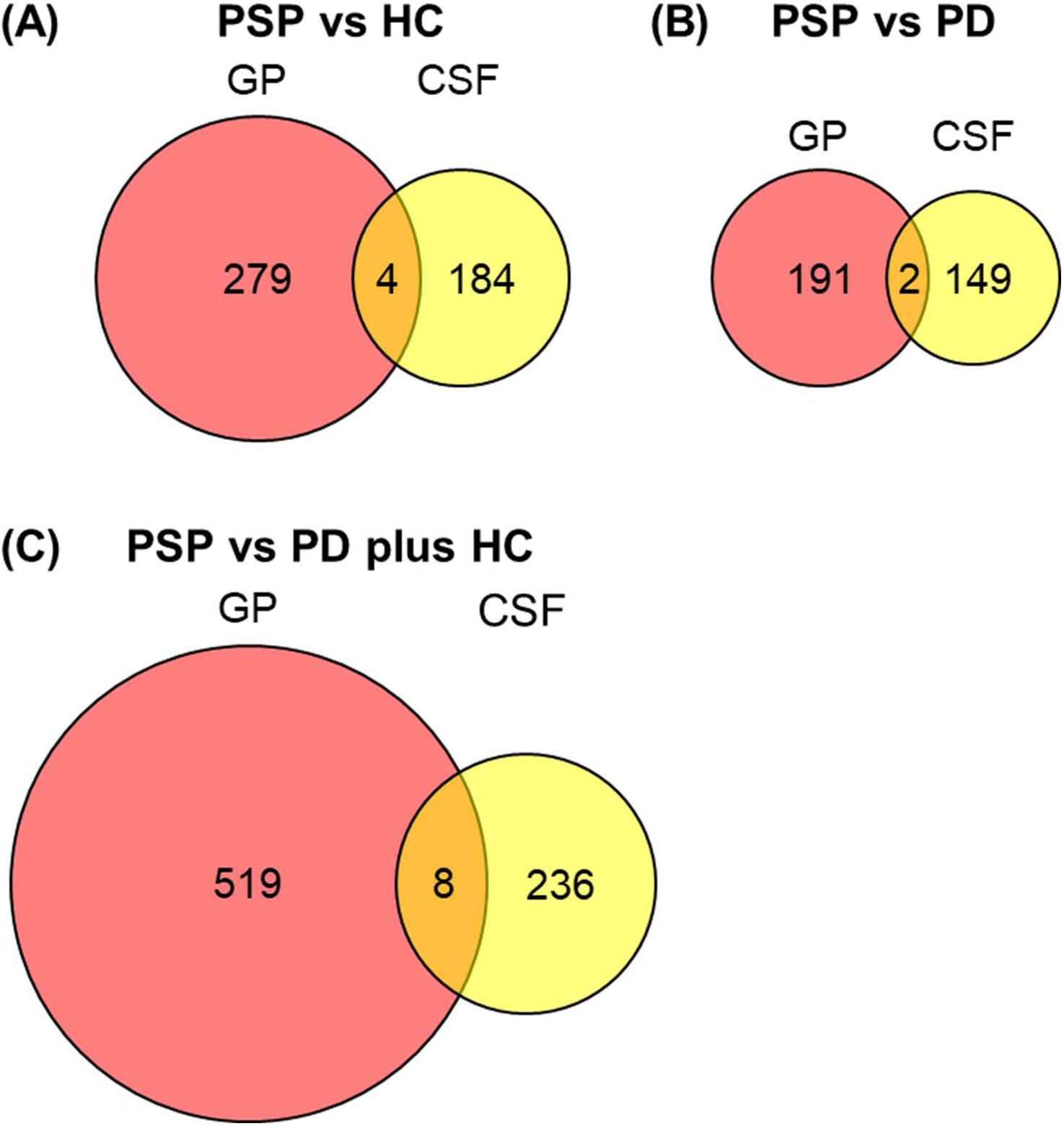
Comparison of differentially expressed proteins in CSF of PSP with the ones in GP of PSP. The differentially expressed proteins in CSF of PSP discovered in this study were compared with the ones in GP of PSP discovered in the previous study.^17^ We used *q*-values < 0.01 for the cutoff to determine differentially expressed proteins.

**Figure 5.**
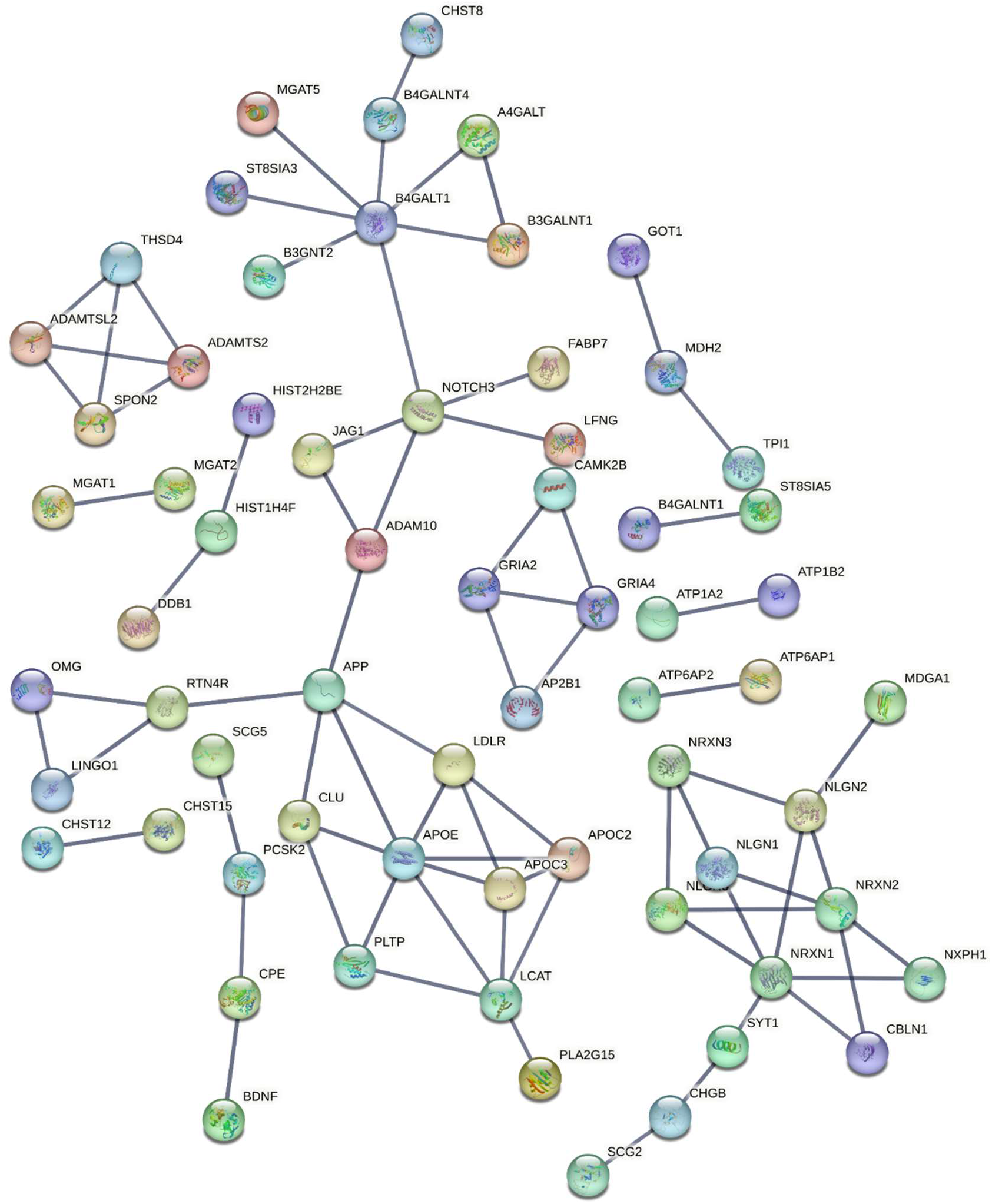
Interactome analysis of differentially expressed proteins. STRING PPI analysis was conducted to estimate the connectivity of the differentially expressed proteins with q-value < 0.01 in PSP compared to the group of HC plus PD. All active interaction sources, including text mining, experiments, databases, co-expression, neighborhood, gene fusion, and co-occurrence, were used with a 0.9 of highest confidence threshold as a minimum required interaction score. Network edges were set to confidence, which indicates data strength based on thickness. The network contains 241 nodes with 76 edges. (average node degree: 0.63, average local clustering coefficient: 0.178, and PPI enrichment *P* value < 1×10^-16^). We selected to hide disconnected nodes in the network.

**Table 2.** Overlapping differentially expressed proteins between PSP GP and PSP CSF.

| PSP vs HC | PSP vs PD | PSP vs PD plus HC |
| --- | --- | --- |
|  |  | ATP1B2 |
| CNTNAP2 |  | CNTNAP2 |
| EPDR1 |  | EPDR1 |
|  |  | FBLN2 |
| GGH | GGH | GGH |
|  |  | GOT1 |
|  | HAPLN4 | HAPLN4 |
|  |  | PREP |
| SERPINE2 |  |  |

### Characterization of differentially expressed proteins in CSF from PSP patients

To better understand the differentially expressed proteins in PSP CSF, we evaluated implicated pathways by gene set enrichment analysis (GSEA). When PSP was compared to HC, the axonal guidance pathway was the most enriched, followed by lysosome pathway, metabolic pathway, cell adhesion molecules pathway, and glycosphingolipid biosynthesis pathway. When PSP was compared to PD, the cell adhesion molecules pathway was the most enriched, followed by cholesterol metabolism pathway, glycosphingolipid biosynthesis pathway, and glycosaminoglycan biosynthesis pathway. When PSP was compared to the group of PD plus HC, cell adhesion molecules pathway was the most enriched, followed by axonal guidance pathway, cholesterol metabolism pathway, lysosome pathway, and various types of N-glycan biosynthesis pathway (Table 3). As expected, proteins known to be implicated in neurodegeneration represent key components in the enriched pathways. Surprisingly, lipid-related proteins were also frequently observed, suggesting their potential connection to the pathogenesis process of PSP.

**Table 3.** KEGG pathway analysis for the differentially expressed proteins.

|  | Term | Count | % | P-Value | Benjamini |
| --- | --- | --- | --- | --- | --- |
| PSP vs<br>HC | Axon guidance | 12 | 6.6 | 8.E-06 | 1.30E-03 |
|  | Lysosome | 8 | 4.4 | 9.E-04 | 6.80E-02 |
|  | Metabolic pathways | 31 | 16.9 | 2.E-03 | 9.30E-02 |
|  | Cell adhesion molecules | 8 | 4.4 | 2.E-03 | 9.30E-02 |
|  | Glycosphingolipid biosynthesis - lacto and neolacto series | 4 | 2.2 | 4.E-03 | 1.20E-01 |
|  | Other types of O-glycan biosynthesis | 4 | 2.2 | 2.E-02 | 4.40E-01 |
|  | N-Glycan biosynthesis | 4 | 2.2 | 2.E-02 | 4.40E-01 |
|  | Glycosaminoglycan biosynthesis - chondroitin sulfate / dermatan sulfate | 3 | 1.6 | 2.E-02 | 4.40E-01 |
|  | Renin-angiotensin system | 3 | 1.6 | 3.E-02 | 5.00E-01 |
|  | Glycosaminoglycan biosynthesis - heparan sulfate / heparin | 3 | 1.6 | 3.E-02 | 5.00E-01 |
|  | Amphetamine addiction | 4 | 2.2 | 5.E-02 | 6.70E-01 |
|  | Various types of N-glycan biosynthesis | 3 | 1.6 | 8.E-02 | 1.00E+00 |
|  | Type I diabetes mellitus | 3 | 1.6 | 9.E-02 | 1.00E+00 |
| PSP vs<br>PD | Cell adhesion molecules | 8 | 5.4 | 4.E-04 | 5.60E-02 |
|  | Cholesterol metabolism | 5 | 3.4 | 1.E-03 | 6.60E-02 |
|  | Glycosphingolipid biosynthesis - lacto and neolacto series | 4 | 2.7 | 2.E-03 | 7.20E-02 |
|  | Glycosaminoglycan biosynthesis - keratan sulfate | 3 | 2 | 7.E-03 | 2.10E-01 |
|  | Other types of O-glycan biosynthesis | 4 | 2.7 | 8.E-03 | 2.10E-01 |
|  | Metabolic pathways | 23 | 15.5 | 1.E-02 | 2.30E-01 |
|  | Glycosaminoglycan biosynthesis - chondroitin sulfate / dermatan sulfate | 3 | 2 | 1.E-02 | 2.50E-01 |
|  | Glycosaminoglycan biosynthesis - heparan sulfate / heparin | 3 | 2 | 2.E-02 | 3.10E-01 |
|  | Lysosome | 5 | 3.4 | 3.E-02 | 4.20E-01 |
|  | Various types of N-glycan biosynthesis | 3 | 2 | 5.E-02 | 6.00E-01 |
|  | Prostate cancer | 4 | 2.7 | 5.E-02 | 6.40E-01 |
|  | N-Glycan biosynthesis | 3 | 2 | 7.E-02 | 7.80E-01 |
|  | Sphingolipid metabolism | 3 | 2 | 8.E-02 | 8.00E-01 |
| PSP vs<br>PD+HC | Cell adhesion molecules | 16 | 6.6 | 6.70E-09 | 1.20E-06 |
|  | Axon guidance | 14 | 5.8 | 2.10E-06 | 1.90E-04 |
|  | Cholesterol metabolism | 7 | 2.9 | 8.80E-05 | 5.30E-03 |
|  | Lysosome | 9 | 3.7 | 6.50E-04 | 2.90E-02 |
|  | Various types of N-glycan biosynthesis | 5 | 2.1 | 3.10E-03 | 1.10E-01 |
|  | Other types of O-glycan biosynthesis | 5 | 2.1 | 4.70E-03 | 1.30E-01 |
|  | Glycosaminoglycan biosynthesis - heparan sulfate / heparin | 4 | 1.7 | 4.80E-03 | 1.30E-01 |
|  | Glycosphingolipid biosynthesis - lacto and neolacto series | 4 | 1.7 | 6.80E-03 | 1.50E-01 |
|  | N-Glycan biosynthesis | 5 | 2.1 | 7.20E-03 | 1.50E-01 |
|  | Metabolic pathways | 33 | 13.7 | 1.90E-02 | 3.50E-01 |
|  | Glycosaminoglycan biosynthesis - chondroitin sulfate / dermatan sulfate | 3 | 1.7 | 3.70E-02 | 6.10E-01 |
|  | Renin-angiotensin system | 3 | 1.2 | 4.30E-02 | 6.60E-01 |
|  | Amphetamine addiction | 4 | 1.7 | 7.90E-02 | 1.00E+00 |
|  | Mucin type O-glycan biosynthesis | 3 | 1.2 | 9.60E-02 | 1.00E+00 |

A protein-protein interaction analysis was conducted. APOE and B4GALT1 were clustered with 7 other proteins, NRXN1 was clustered with 6 other proteins, and APP, LCAT, NOTCH3 and NRXN2 were clustered with 5 other proteins. This interaction analysis suggested that APOE, B4GALT1, NRXN1, NRXN2, APP, LCAT, and NOTCH3 were key components. Among them, APOE, B4GALT1, NRXN1, NRXN2, and LCAT are involved in the cell adhesion molecules pathway, cholesterol metabolism pathway, and glycan biosynthesis pathway in PSP vs PD plus HC of GSEA, further suggesting a potential connection of cell adhesion molecule and cholesterol metabolism pathways to PSP.

Cell-type-specific enrichment analysis was performed to characterize the differentially expressed proteins in PSP compared to the group of PD plus HC. Neuronal proteins were the most enriched when the top 50 up and down regulated proteins were analyzed, followed by astrocytic proteins. Neuronal proteins were also the most enriched when examining all differentially expressed proteins, while oligodendrocytic proteins were the second most enriched (Table 4). Neuronal proteinsdetected in CSF are the primary group of proteins that change followed by oligodendrocytic proteins in PSP probably reflecting loss of neurons in PSP.

**Table 4.** Cell-type-specific enrichment of proteins differential between PSP and PD plus HC.

|  | Cell type | *P* value | List of differential proteins that overlap with the proteins enriched in a specific cell type |
| --- | --- | --- | --- |
| Top 50 up- and down-regulated proteins with *q*-value < 0.01 | Astrocyte | 0.039 | LCAT, TIMP3, CRYM, BTD, SLITRK2, MFGE8, BDNF, PDYN |
|  | Microglia | 0.093 | LAMP2, CHST12, B4GALT1, ADAM15, GGH, KCTD12, GBA |
|  | Neurons | 0.015 | NDRG4, MINPP1, ST8SIA5, CDH7, CLSTN3, CHGB, FAT2 |
|  | Oligodendrocyte | 0.353 | PCSK1N, LINGO3, SLITRK4, ADAMTS2, SERPINA3 |
|  | Endothelia | 0.556 | B3GNT2, A4GALT, SEMA3G, PREP |
| All the differential proteins with *q*-value < 0.01 | Astrocyte | 0.026 | PRDX6, BTD, ALDH1A1, FGFR1, BDNF, MFGE8, PDYN, TRIL, TIMP3, SLITRK2, CRYM, LCAT, ATP1A2, ATP1B2, SCG3, CPE, LINGO1, NOTCH3, NRXN1, APOE, LRIG1, CLU, FABP7, VNN1, NRCAM, EPHB3, EDNRB |
|  | Microglia | 0.026 | GBA, ASPH, OLFML3, KCTD12, APOC2, ADAM15, GGH, B4GALT1, CHST12, LAMP2, HPRT1, HS6ST1, DPP7, AP1B1, ADGRB1, MGAT2, EPDR1, CORO1A, B4GALNT1, CDH23, LFNG, FAM3A, PLA2G15, MGAT1, PLOD1, CTSA, LIPA |
|  | Neurons | 4.68E-11 | TRHDE, XXYL1, NRSN2, ST8SIA5, CLSTN3, CBLN1, ST8SIA3, MINPP1, MGAT5, CDH7, CHGB, NDRG4, FAT2, NPTX1, SCN4B, NRN1, CNTNAP2, EPHA6, PCDHAC2, TMEM59L, NXPH3, SEMA6C, ADGRL1, CBLN4, SCG2, SERPINI1, VGF, L1CAM, RTN4R, RGMB, SCN2B, CA10, CNTNAP5, CAMK2B, MMP24, ROBO2, IDS, CNTNAP4, CACNA2D2, SYT1, CLSTN1, FSTL4, PCSK1, DNER, VWC2, PENK, B4GALNT4, MDGA1, PCDH17, PTPRN2 |
|  | Oligodendrocyte | 0.009 | QPCT, SLITRK4, ADAMTS2, GALNT13, LINGO3, EXTL2, SERPINA3, PCSK1N, PCDH7, NFASC, SEMA6A, NPTX2, TMEM132C, SEMA4D, GPR158, NLGN3, ADAM11, ADGRL3, FBLN7, TMEM132D, MFAP4, CA10, LDLR, XYLT1, CHST8, NXPH1, SERPINE2, BRINP1, LRFN2 |
|  | Endothelia | 0.280 | SPOCK2, PREP, SLC39A10, MMRN2, SEMA3G, A4GALT, B3GNT2, HPRT1, GINM1, TFRC, JAG1, ITM2A, ADAMTS2, PTPRM, SEMA7A, SPON2, FBLN5, GALNT18, PLTP, ART3, ADAM10 |

### Evaluation of the candidate biomarker proteins for classification performance

As the main goal of this study was to identify proteins that can be used to differentiate PSP from HC and PD, we evaluated the classification performance of differentially expressed proteins using ROC analysis. ATP6AP2 showed the highest AUC value (0.922), followed by NEFM (AUC 0.894), EFEMP2 (AUC 0.892), LAMP2 (AUC 0.845), CHST12 (AUC 0.838), FAT2 (AUC 0.810), B4GALT1 (AUC 0.808), LCAT (AUC 0.800), CBLN3 (AUC 0.792), FSTL5 (AUC 0.791), ATP6AP1 (AUC 0.790), and GGH (AUC 0.789) (Figure 6). The remainder of the top 50 up- and down-regulated proteins showed AUC > 0.696 (Supplemental Figure S1). To further improve the classification performance of differentially expressed proteins, we conducted multivariate ROC analyses by varying the number of features. When 53 features were used, they showed the highest AUC (0.937). However, the classification performance using the top 3 features did not show significant difference from the one using 53 features, suggesting that 3 main proteins contributed to the majority of classification performance (Figure 7A). Using the 53 features, we conducted 3D PCA analysis, and most PSP samples showed clear separation from PD and HC with few exceptions (Figure 7B). We next evaluated the predictive accuracy of the features. When 5 features were used, it reached 94.1% (Figure 3C). We evaluated what proteins contribute the most to the classification performance. NEFM contributed the most followed by CHI3L1, ATP6AP2, LAMP2, CHGB, GRIA4, GGH, FAT2, ENPP5, BDNF, CBLN3, SERPINE2, ZP2, CDH7, and FSTL5. The relative abundances of the top 53 proteins are shown in Supplemental Figure S2. These data suggest that the combination of these potential biomarker proteins can be used to successfully differentiate PSP from PD and HC individuals.

**Figure 6.**
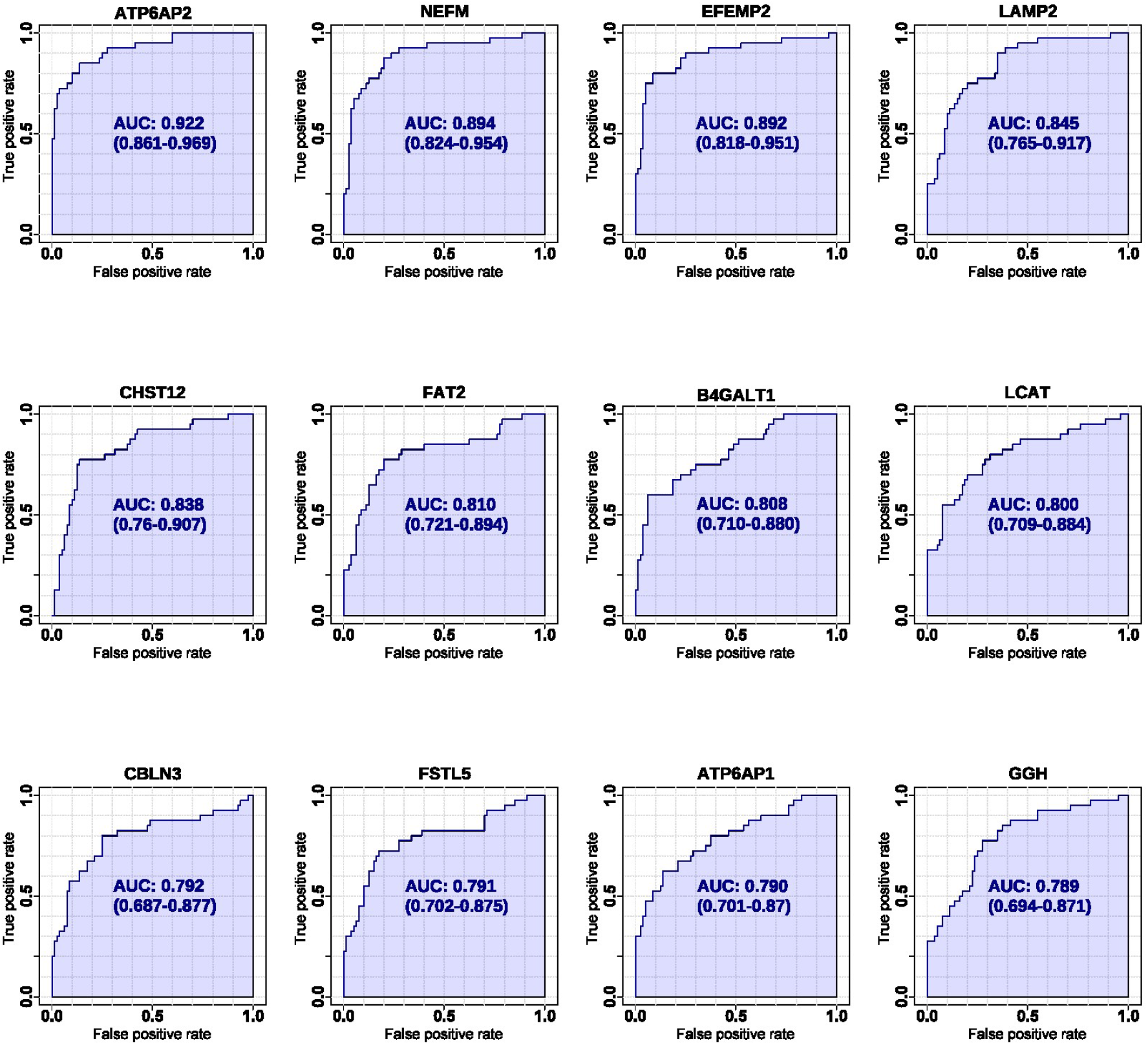
ROC analysis of 12 representative proteins with the highest AUCs between PSP vs PD plus HC. ROC analysis was generated by bootstrapping. The values in the parenthesis show the lower and upper AUC values of 95% confidence interval.

**Figure 7.**
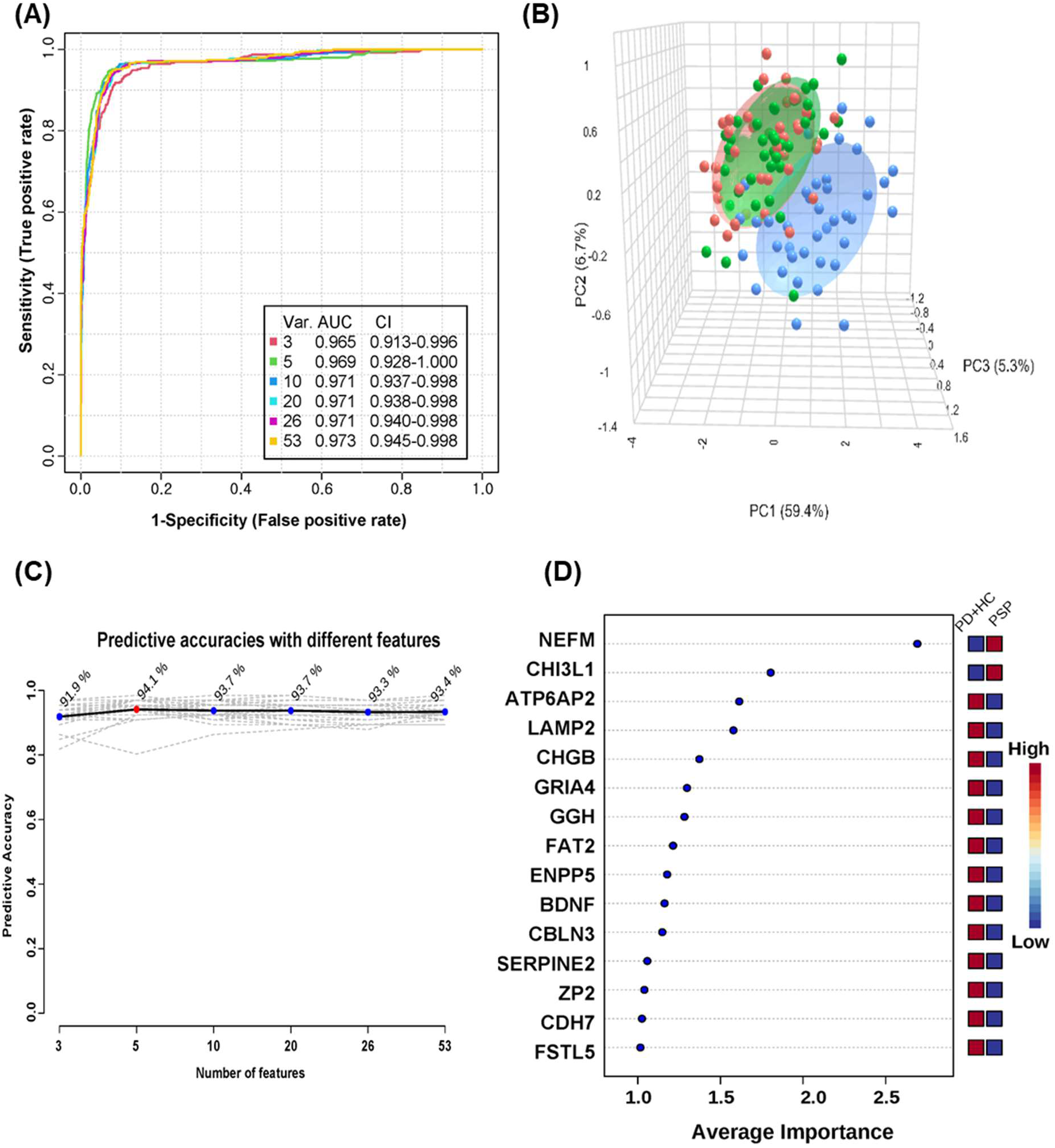
Multivariate ROC analysis and predictive accuracy. The differentially expressed proteins of PSP-specific biomarker candidates were compared with HC plus PD. Multivariate ROC analyses were conducted using different numbers of features ranging from 3 to 53. Var. indicates the number of features used. CI indicates confidence interval (A). Principal Component Analysis for top 53 differential proteins between PSP vs PD plus HC (B). The accuracy for predicting PSP as the number of features increased is shown (C). The top 15 significant features affecting the discrimination of PSP from PD plus HC are shown with their average importance (D).

## DISCUSSION

In this study, mass spectrometry-based proteomic analysis of 120 human CSF samples from 40 PSP, 40 PD, and 40 HC individuals was conducted using the TMT-based multiplexing approach, identifying 3,653 proteins. Although we analyzed 120 CSF samples using 13 batches of 11-plex TMT experiments, the precision of the experiment was very high, with < 10% of CV for most of the proteins. This suggests that the two-step normalization using MP and, subsequently, ComBat package was effective for the analysis of large number of CSF samples using the TMT-based quantification approach.

Since we wanted to explore whether the differentially expressed proteins in both CSF and GP of PSP patients may have greater relevance as PSP biomarkers, we built on our prior work and compared the proteins differentially expressed in both CSF and GP. While ATP1B2, CNTNAP2, EPDR1, FBLN2, GGH, GOT1, HAPLN4, PREP, and SERPINE2 were the main differentially expressed proteins common in CSF and GP of PSP patients, they were not identified as the key proteins in GSEA, interactome analysis, and ROC analyses. This discrepancy suggests that GP-derived proteins may not reflect the same pathogenic process giving rise to differentially expressed proteins in PSP CSF. Rather, because multiple other brain regions (such as the subthalamic nucleus, substantia nigra, putamen and perirolandic cortex) are prominently affected by PSP pathology, evaluating protein expression in these additional regions may yield results in greater concordance with the CSF findings.^28,29^ Furthermore, because autopsy samples are mostly derived from patients in advanced stages of PSP, the differentially expressed proteins in early or mid-stage PSP could well be different from the ones expressed in the advanced stages—reflecting a dynamic pathophysiological process. Further investigations utilizing CSF samples derived from multiple disease stages and autopsy samples derived from multiple implicated brain regions are necessary to further characterize the proteomic signature of PSP.

GSEA and interactome analysis demonstrated that cell adhesion molecules pathway, cholesterol metabolism pathway, and glycan biosynthesis pathway were the critical ones for the differentially expressed proteins in PSP CSFs. In these pathways, APOE, B4GALT1, NRXN1, NRXN2, and LCAT were key proteins. Cell adhesion molecules are already known to be involved in neurodegenerative diseases,^30,31^ especially by altering synaptic plasticity, neuroinflammatory events, and effecting vascular changes. Cholesterol is an indispensable component of the cell membrane, and aberrations of cholesterol metabolism are involved in various neurodegenerative conditions, including Alzheimer’s disease (AD) and PD.^32^ Glycan is a key molecule involved in the modification of lipids, proteins and other glycans.^33^ Glycosylated lipids are involved in cell adhesion and glycosylated proteins are major components of cell membrane proteins.^34^ Our cell-type enrichment analysis results indicated that the main fraction of the differentially expressed proteins was derived from neuronal cells, suggesting that the pathway changes observed in CSF were predominantly from neuronal cells; further investigation is required to validate this.

The primary aim of this study was to identify biomarkers for PSP. To this end, we assessed the discriminatory potential of several candidate biomarker proteins for PSP. ATP6AP2 had the highest AUC, followed by NEFM, EFEMP2, LAMP2, CHST12, FAT2, B4GALT1, LCAT, CBLN3, FSTL5, ATP6AP1, and GGH, when compared to PD and HC. Of the top 12 proteins, B4GALT1 plays a role in glycan biosynthesis, while LCAT is involved in cholesterol metabolism. Both these proteins emerged as significant in our interactome analysis as well, suggesting that B4GALT1 and LCAT might be promising novel biomarkers for PSP.

B4GALT1 is a galactosyltransferase enzyme, which is responsible for the synthesis of oligosaccharides in glycoproteins and glycolipids. B4GALT1 is known to be linked to microglial activation and neuroinflammation.^35^ In AD brains, elevated B4GALT1 expression correlated with heightened galactosylation of N-glycans.^36^ Furthermore, a previous study indicated a notable increase in B4GALT1 gene expression within the substantia nigra of PD patients compared to controls.^35^ However, our result showed downregulation of B4GALT1 in the CSF from PSP patients when compared to PD and HC. Further investigation is required to assess whether and how B4GALT1 is involved in the pathogenesis of PSP.

LCAT is a lipoprotein-associated enzyme that plays a key role in transferring excessive cholesterol in peripheral tissues to the liver for excretion.^37,38^ The dysregulation of LCAT leads to the disturbance of lipid metabolism and it is potentially implicated in the pathogenesis of PD.^39^ Recent plasma metabolomics analyses underscore this by revealing a decrease in lipid and lipid-associated molecules in PD compared to the control group.^40^ Our findings showed that LCAT was downregulated in PSP patients compared to PD and HC, suggesting the link between lipid metabolism disturbance by LCAT dysregulation and PSP pathogenesis.

ATP6AP2 is ATPase H+ transporting lysosomal accessory protein, which is a vital component of the vacuolar ATPase and plays a crucial role in lysosomal functions and autophagy. Deficiency in ATP6AP2 disrupts V-ATPase function, affecting neural stem cell renewal and causing widespread neural degeneration, emphasizing ATP6AP2’s central role in the developing human nervous system.^41^ The dysregulation of ATP6AP2 was also reported to be implicated in Parkinsonism.^42^ Our findings indicate a decreased expression of ATP6AP2 in PSP patients relative to both HC and PD, suggesting that ATP6AP2 dysregulation plays a role in PSP. Notably, the mode of ATP6AP2 dysregulation in PSP appears distinct from that in PD, given the differential levels observed between the two patient groups. Further study is required to investigate this distinction.

Neurofilament proteins, including NEFM, are considered promising candidate state biomarkers for neuronal damage and the process of neurodegeneration.^43^ However, they are relatively non-specific when attempting to differentiate among neurological diseases diagnostically. Elevated levels within CSF have been demonstrated for patients with stroke and a wide spectrum of neurodegenerative and neuroinflammatory conditions.^44,45^ While there is a lack of research on the relationship between neurofilament proteins and PSP, our findings suggest that PSP patients sustain significant ongoing neuronal damage and thus release greater amounts of NEFM into CSF compared to PD and HC individuals.

EFEMP2, also known as fibulin-4, is a member of the fibulin glycoprotein family found predominantly in elastic fiber-rich tissues and is vital for elastic fiber formation, connective tissue development, and extracellular matrix stability.^46^ EFEMP2 has been reported to have implications in the advancement of different cancer types.^47^ Little is known about the relationship between EFEMP2 and neurodegeneration. We found downregulation of EFEMP2 in PSP patients compared to HC and PD, and further investigation of this relationship is required.

LAMP2 is a lysosomal-associated membrane protein and constitutes a significant portion of the lysosomal membrane.^48^ Lysosomes serve as the main catabolic units responsible for breaking down intracellular proteins via the process of autophagy.^49^ The existence of α-synuclein aggregates in PD is potentially mediated by compromised degradation capabilities of lysosomes.^50^ A prior investigation using Western blot quantification reported that PD CSFs showed reduced levels of LAMP2 compared to HC, while PSP CSF did not show differences.^51^ Our result showed a downregulation of LAMP2 in PSP patients compared to HC and PD. This discrepancy could be caused by quantification method differences or case specificity, and further investigation is required to clarify this.

CHST12 is a carbohydrate sulfotransferase involved in the biosynthesis of proteoglycans that facilitate cell interactions. Its overexpression serves as an unfavorable prognostic factor in ovarian cancer.^52^ Little is known about the involvement of CHST12 in neurodegeneration. We found that the CHST12 level was decreased in PSP compared to PD and HC.

FAT2 is a cadherin superfamily protein and is known to be expressed in granule cells in the cerebellum.^53^ The cadherin family proteins have consistently demonstrated their influence in governing the contact between axons and dendrites.^54^ Our results showed a downregulation of FAT2 in PSP patients compared to PD and HC. Further investigation is required to understand how FAT2 is involved in PSP.

CBLN3 is a member of the precerebellin protein family^55^ and is expressed in cerebellum and dorsal cochlear nucleus.^55^ The link between CBLN3 and neurodegeneration is not clear, although our finding shows that CBLN3 was downregulated in PSP compared to PD and HC and cerebellar pathology (particularly in the dentate) is well-described in PSP.^56,57^

FSTL5 is a secretory glycoprotein^58^ and is known to be a prognostic biomarker for medulloblastoma.^59^ Our data showed that FSTL5 was significantly downregulated in PSP compared PD and HC.

ATP6AP1 is an accessory protein of V-type ATPase proton pump.^60^ Its role is to direct the V-ATPase to specific subcellular compartments, such as neuroendocrine-regulated secretory vesicles, and to regulate various aspects of their function, including intragranular pH and the Ca2+-dependent exocytotic membrane fusion.^46^ In our results, ATP6AP1 showed significant downregulation in the PSP compared to PD and HC. Considering that both ATP6AP1 and ATP6AP2 are downregulated in PSP, the subcellular mislocalization of V-type ATPase proton pump by the dysfunction of its accessory proteins is potentially involved in PSP pathogenesis.

GGH is an enzyme involved in folate metabolism.^61^ Fang *et al.* reported GGH was downregulated in human CSF from Huntington disease patients.^62^ Licker *et al.* also reported that GGH was downregulated in substantia nigra of PD patients.^63^ Our finding also showed that GGH was significantly downregulated in PSP compared to PD and HC. These studies suggest that GGH is downregulated in multiple various neurodegenerative diseases.

In this study, NPTX2 was downregulated in PSP compared to PD and HC. The downregulation of NPTX2 is a predictive marker for the progression from normal cognition to mild cognitive impairment,^25^ and cognitive decline is a typical symptom of PSP.^64^ This suggests that dysregulated synaptic adaptability mediated by NPTX2 downregulation could be a potential mechanism of the cognitive decline of PSP patients.

Multivariate analysis showed marginally improved discriminating capability (AUC 0.937) compared to the best single-marker AUC (0.922) of ATP6AP2. This suggests that ATP6AP2 is a promising single-marker candidate for PSP and that integrating multiple PSP biomarkers could be beneficial for the better diagnosis of PSP.

Important study limitations include the lack of post-mortem confirmation of PSP or PD diagnosis and differences between groups with respect to age, race/ethnicity and education. Every effort to match samples on demographic characteristics was made, but we acknowledge that these differences may have contributed to differential CSF protein expression in ways that are not currently well understood. It should be noted that lower education levels have previously been associated with higher likelihood of a PSP diagnosis,^65^ though the pathophysiological mechanism of this association remains unclear. The candidate biomarkers discovered in this study also need to be validated using an independent cohort and also evaluated for their applicability to differentiate across subtypes of PSP.

To the best of our knowledge, this is the first global-scale proteome analysis to discover CSF PSP biomarkers using a mass spectrometry-based proteomics approach and utilizing samples from well-matched PSP, PD, and controls. This study provides the foundation for the development of PSP biomarkers, which should be validated in multi-center cohorts.

## ACKNOWLEDGMENTS

We acknowledge an NIH shared instrumentation grant (S10OD021844 to T.M.D.). T.M.D is the Leonard and Madlyn Abramson Professor in Neurodegenerative Diseases.

## FUNDING

This work was supported by an NIH grant (U01 NS102035 to A.Y. P and T.M.D.).

## CONFLICTS OF INTERESTS

We have no conflict of interest to declare.

## AUTHOR CONTRIBUTIONS

Z.Z., T.M.D., C.H.N., and A.Y.P. designed research; A.J.H., T.F.T., and A.C.-P prepared and shared CSF samples; Y.J. conducted the mass spectrometry experiments; Y.J., S.O., Z.Z., and C.H.N. performed data analysis; Y.J, S.O., Z.Z., T.F.T, A.C.-P., L.S.R., T.M.D., C.H.N., and A.Y.P. wrote the manuscript; T.M.D., C.H.N., and A.Y.P. supervised research.

## ETHICAL APPROVAL

This research study was approved by the Johns Hopkins Institutional Review Board (IRB Application #00173663). This study abided by the Declaration of Helsinki principles.

## Abbreviations

ACN: acetonitrile
AGC: automatic gain control
AUC: area under the curve
bRPLC: basic pH reversed-phase liquid chromatography
CAA: chloroacetamide
CSF: cerebrospinal fluid
CV: coefficient variation
DDA: data-dependent acquisition
FA: formic acid
FDR: false-discovery rate
HC: healthy control
HCD: higher-energy collisional dissociation
KEGG: Kyoto encyclopedia of genes and genomes
m/z: mass-to-charge ratio
MP: master pool
PD: Parkinson’s disease
PSM: peptide-spectrum match
PSP: progressive supranuclear palsy
QC: quality control
ROC: receiver operating characteristic
RT: room temperature
S/N: signal-to-noise ratios
SD: standard deviation
TCEP: tris (2-Carboxyethyl) phosphine hydrochloride
TEAB: triethylammonium bicarbonate
TFA: trifluoroacetic acid
TMT: tandem mass tag

---

**Supplemental Table S1.** Top 50 down- and up-regulated differential proteins in PSP compared to HC.

| No. | Protein name | P value | Mean of bootstrap AUC | SD of bootstrap AUC | q-value |
| --- | --- | --- | --- | --- | --- |
| Down-regulated proteins |  |  |  |  |  |
| 1 | ATP6AP2 | 4.97E-14 | 0.07375 | 0.0285685 | 0 |
| 2 | CHST12 | 2.66E-08 | 0.156875 | 0.0435088 | 0 |
| 3 | EFEMP2 | 4.46E-10 | 0.126875 | 0.0443257 | 0 |
| 4 | LAMP2 | 1.34E-07 | 0.18125 | 0.0468534 | 0 |
| 5 | ATP6AP1 | 5.35E-08 | 0.176875 | 0.0470242 | 0 |
| 6 | PCMT1 | 4.86E-07 | 0.19375 | 0.0474892 | 0 |
| 7 | NDRG4 | 2.96E-06 | 0.20875 | 0.0480604 | 0 |
| 8 | FSTL5 | 1.92E-07 | 0.178125 | 0.0480624 | 0 |
| 9 | FAT2 | 6.77E-07 | 0.194375 | 0.0481374 | 0 |
| 10 | FBLN2 | 9.20E-06 | 0.22125 | 0.0488661 | 0 |
| 11 | CBLN3 | 2.82E-06 | 0.20625 | 0.0489301 | 0 |
| 12 | PCSK2 | 3.35E-06 | 0.20875 | 0.0489551 | 0 |
| 13 | ZP2 | 1.84E-06 | 0.201875 | 0.0490643 | 0 |
| 14 | B3GNT2 | 4.02E-06 | 0.22125 | 0.0499495 | 0 |
| 15 | CLSTN3 | 2.69E-06 | 0.21625 | 0.0506021 | 0 |
| 16 | SEMA3G | 1.05E-05 | 0.224375 | 0.0506108 | 0 |
| 17 | CDH7 | 6.47E-06 | 0.229375 | 0.0507745 | 0 |
| 18 | ANGPTL2 | 3.70E-06 | 0.218125 | 0.0509816 | 0 |
| 19 | B4GALT1 | 6.02E-06 | 0.224375 | 0.0513025 | 0 |
| 20 | MFGE8 | 9.75E-06 | 0.22375 | 0.0514011 | 0 |
| 21 | PREP | 5.02E-06 | 0.21375 | 0.0515128 | 0 |
| 22 | LINGO3 | 1.15E-05 | 0.229375 | 0.0517718 | 0 |
| 23 | PCSK1N | 1.27E-06 | 0.205625 | 0.0517753 | 0 |
| 24 | BDNF | 5.46E-06 | 0.22125 | 0.0520363 | 0 |
| 25 | GGH | 2.33E-05 | 0.23125 | 0.0521141 | 0 |
| 26 | GOT1 | 4.12E-05 | 0.25375 | 0.0521524 | 0 |
| 27 | ACP2 | 3.69E-05 | 0.23625 | 0.0522678 | 0 |
| 28 | LCAT | 9.67E-06 | 0.229375 | 0.0523143 | 0 |
| 29 | ENPP5 | 1.45E-05 | 0.2225 | 0.0523766 | 0 |
| 30 | OLFML3 | 1.67E-05 | 0.2225 | 0.0524193 | 0 |
| 31 | ADAM15 | 3.02E-05 | 0.2384375 | 0.0524369 | 0 |
| 32 | MGAT5 | 5.03E-06 | 0.215625 | 0.0525166 | 0 |
| 33 | TPI1 | 2.45E-05 | 0.24375 | 0.0527149 | 0 |
| 34 | CRYM | 9.85E-06 | 0.24125 | 0.0528349 | 0 |
| 35 | EXTL2 | 3.50E-06 | 0.225625 | 0.052838 | 0 |
| 36 | XXYL1 | 3.70E-05 | 0.239375 | 0.0530386 | 0 |
| 37 | QSOX2 | 9.12E-05 | 0.244375 | 0.0533785 | 0 |
| 38 | GALNT13 | 4.72E-05 | 0.24375 | 0.053383 | 0 |
| 39 | ST8SIA3 | 3.56E-06 | 0.22625 | 0.0534163 | 0 |
| 40 | FAM3A | 3.42E-05 | 0.24875 | 0.0534198 | 0 |
| 41 | MINPP1 | 5.10E-05 | 0.251875 | 0.0534487 | 0 |
| 42 | CHGB | 1.08E-05 | 0.2325 | 0.0536 | 0 |
| 43 | NUTF2 | 3.65E-05 | 0.25 | 0.0536198 | 0 |
| 44 | EPHA6 | 3.76E-05 | 0.25125 | 0.0536506 | 0 |
| 45 | CA11 | 8.76E-06 | 0.228125 | 0.0538124 | 0 |
| 46 | SRPX | 8.47E-05 | 0.244375 | 0.053815 | 0 |
| 47 | TIMP4 | 1.37E-05 | 0.234375 | 0.053927 | 0 |
| 48 | ARL8B | 1.57E-05 | 0.238125 | 0.054094 | 0 |
| 49 | SPOCK2 | 1.76E-05 | 0.2425 | 0.054095 | 0 |
| 50 | SLC39A10 | 7.67E-05 | 0.259375 | 0.0541175 | 0 |
| Up-regulated proteins |  |  |  |  |  |
| 1 | NEFM | 8.17E-10 | 0.8675 | 0.0432413 | 0 |
| 2 | CHI3L1 | 5.45E-05 | 0.753125 | 0.053142 | 0 |
| 3 | SERPINA3 | 2.40E-05 | 0.76 | 0.0539946 | 0 |
| 4 | MMRN1 | 0.0010768 | 0.726875 | 0.0563351 | 0 |
| 5 | SELL | 0.0023944 | 0.7025 | 0.0599897 | 0 |

**Supplemental Table S2.** Top 50 down- and up-regulated differential proteins in PSP compared to PD.

| No. | Protein name | P value | Mean of bootstrap AUC | SD of bootstrap AUC | q-value |
| --- | --- | --- | --- | --- | --- |
| Down-regulated proteins |  |  |  |  |  |
| 1 | ATP6AP2 | 1.19E-13 | 0.081875 | 0.0320833 | 0 |
| 2 | EFEMP2 | 1.92E-11 | 0.088125 | 0.0348503 | 0 |
| 3 | LAMP2 | 2.61E-10 | 0.131875 | 0.0410988 | 0 |
| 4 | B4GALT1 | 1.46E-08 | 0.165 | 0.0459937 | 0 |
| 5 | GGH | 9.38E-07 | 0.19125 | 0.0478408 | 0 |
| 6 | APOC2 | 3.88E-06 | 0.204375 | 0.0489066 | 0 |
| 7 | CHST12 | 4.74E-07 | 0.1725 | 0.0490566 | 0 |
| 8 | LCAT | 1.87E-07 | 0.17375 | 0.0493278 | 0 |
| 9 | FBLN2 | 8.40E-06 | 0.229375 | 0.0511735 | 0 |
| 10 | MINPP1 | 2.12E-05 | 0.24 | 0.0517964 | 0 |
| 11 | CHGB | 7.93E-05 | 0.255625 | 0.0518278 | 0 |
| 12 | APOC3 | 5.12E-05 | 0.243125 | 0.0520782 | 0 |
| 13 | ADAM15 | 1.95E-05 | 0.234375 | 0.0520889 | 0 |
| 14 | PCMT1 | 3.14E-05 | 0.249375 | 0.0521856 | 0 |
| 15 | FGFR1 | 3.59E-05 | 0.244375 | 0.0521913 | 0 |
| 16 | CBLN3 | 5.35E-06 | 0.211875 | 0.0522269 | 0 |
| 17 | THSD4 | 8.84E-06 | 0.2275 | 0.0522725 | 0 |
| 18 | GRIA4 | 6.15E-05 | 0.240625 | 0.0525096 | 0 |
| 19 | FAT2 | 1.61E-06 | 0.195625 | 0.0528504 | 0 |
| 20 | ST8SIA3 | 0.00013 | 0.26625 | 0.0529591 | 0 |
| 21 | SELENOF | 0.0001655 | 0.254375 | 0.0530813 | 0 |
| 22 | ZP2 | 0.0001129 | 0.255625 | 0.0531251 | 0 |
| 23 | KCTD12 | 2.98E-06 | 0.2225 | 0.0531874 | 0 |
| 24 | ATP6AP1 | 1.58E-05 | 0.243125 | 0.0533337 | 0 |
| 25 | A4GALT | 8.73E-05 | 0.24 | 0.0534313 | 0 |
| 26 | NDRG4 | 3.48E-05 | 0.243125 | 0.054273 | 0 |
| 27 | CRYM | 0.0001416 | 0.264375 | 0.0544155 | 0 |
| 28 | C1QTNF3-AMACR | 1.37E-05 | 0.22375 | 0.0544182 | 0 |
| 29 | CHST10 | 4.91E-05 | 0.255625 | 0.0550191 | 0 |
| 30 | ST8SIA5 | 4.80E-05 | 0.241875 | 0.0550358 | 0 |
| 31 | GBA | 0.0001015 | 0.250625 | 0.0555266 | 0 |
| 32 | BTB | 0.0001037 | 0.258125 | 0.0555498 | 0 |
| 33 | B3GNT2 | 5.41E-05 | 0.26 | 0.0556641 | 0 |
| 34 | ANGPTL2 | 0.0001193 | 0.239375 | 0.0560282 | 0 |
| 35 | CBLN1 | 0.000115 | 0.261875 | 0.0560959 | 0 |
| 36 | METRN | 5.11E-05 | 0.234375 | 0.0561886 | 0 |
| 37 | ASPH | 4.65E-05 | 0.25125 | 0.0562954 | 0 |
| 38 | PCSK1N | 0.0001431 | 0.263125 | 0.0566301 | 0 |
| 39 | CDH7 | 0.0001113 | 0.259375 | 0.0567889 | 0 |
| 40 | SLITRK2 | 9.60E-05 | 0.253125 | 0.0573855 | 0 |
| 41 | FSTL5 | 0.0001379 | 0.245625 | 0.058851 | 0 |
| 42 | NOMO2 | 0.0002514 | 0.27 | 0.0550972 | 0.0067114 |
| 43 | NRXN2 | 0.0002897 | 0.27625 | 0.0554634 | 0.0067114 |
| 44 | MCAM | 0.0002717 | 0.293125 | 0.0554791 | 0.0067114 |
| 45 | GALNT13 | 0.0002 | 0.27625 | 0.0555443 | 0.0067114 |
| 46 | CEMIP | 0.0010199 | 0.300625 | 0.0557235 | 0.0067114 |
| 47 | QSOX2 | 0.0001914 | 0.271875 | 0.0557939 | 0.0067114 |
| 48 | TFRC | 0.0003821 | 0.286875 | 0.0558295 | 0.0067114 |
| 49 | FGFR1 | 0.0003979 | 0.284375 | 0.0559385 | 0.0067114 |
| 50 | TRIL | 0.000499 | 0.281875 | 0.0560925 | 0.0067114 |
| Up-regulated proteins |  |  |  |  |  |
| 1 | NEFM | 2.70E-12 | 0.9175 | 0.0328792 | 0 |
| 2 | SERPINA3 | 1.31E-05 | 0.7675 | 0.0514578 | 0 |
| 3 | CHI3L1 | 7.27E-05 | 0.751875 | 0.05568 | 0 |

**Supplemental Table S3.** Top 50 down- and up-regulated differential proteins in PSP compared to PD plus HC.

| No. | Protein name | P value | Mean of bootstrap AUC | SD of bootstrap AUC | q-value |
| --- | --- | --- | --- | --- | --- |
| Down-regulated proteins |  |  |  |  |  |
| 1 | ATP6AP2 | 2.55E-15 | 0.0778125 | 0.0262204 | 0 |
| 2 | EFEMP2 | 5.43E-12 | 0.1075 | 0.0357351 | 0 |
| 3 | LAMP2 | 2.60E-11 | 0.1565625 | 0.0367626 | 0 |
| 4 | CHST12 | 8.79E-10 | 0.1646875 | 0.0400604 | 0 |
| 5 | NDRG4 | 9.75E-07 | 0.2259375 | 0.0426844 | 0 |
| 6 | B4GALT1 | 2.65E-08 | 0.1946875 | 0.0430197 | 0 |
| 7 | MINPP1 | 4.12E-06 | 0.2459375 | 0.0436766 | 0 |
| 8 | ADAM15 | 6.01E-07 | 0.2364063 | 0.0444755 | 0 |
| 9 | ST8SIA5 | 1.97E-05 | 0.26 | 0.0448256 | 0 |
| 10 | B3GNT2 | 1.24E-06 | 0.240625 | 0.0448994 | 0 |
| 11 | ZP2 | 6.44E-07 | 0.22875 | 0.0451018 | 0 |
| 12 | ANGPTL2 | 1.11E-06 | 0.22875 | 0.0455461 | 0 |
| 13 | LCAT | 2.18E-07 | 0.2015625 | 0.045723 | 0 |
| 14 | GGH | 5.41E-08 | 0.21125 | 0.0457514 | 0 |
| 15 | TIMP3 | 1.29E-05 | 0.2690625 | 0.0458 | 0 |
| 16 | CDH7 | 6.72E-06 | 0.244375 | 0.0459107 | 0 |
| 17 | FBLN2 | 5.84E-07 | 0.2253125 | 0.0460933 | 0 |
| 18 | CRYM | 3.07E-06 | 0.2528125 | 0.0462657 | 0 |
| 19 | ATP6AP1 | 1.06E-07 | 0.21 | 0.0462855 | 0 |
| 20 | PCMT1 | 6.95E-07 | 0.2215625 | 0.0463149 | 0 |
| 21 | CLSTN3 | 1.27E-05 | 0.2584375 | 0.0463512 | 0 |
| 22 | PCSK1N | 2.31E-06 | 0.234375 | 0.0465225 | 0 |
| 23 | LINGO3 | 9.27E-06 | 0.255 | 0.0468393 | 0 |
| 24 | BTD | 2.00E-05 | 0.2775 | 0.0469606 | 0 |
| 25 | ENPP5 | 8.28E-06 | 0.2503125 | 0.0470233 | 0 |
| 26 | CHGB | 7.71E-06 | 0.2440625 | 0.0470607 | 0 |
| 27 | A4GALT | 1.28E-05 | 0.259375 | 0.0470748 | 0 |
| 28 | CBLN3 | 5.19E-07 | 0.2090625 | 0.0472451 | 0 |
| 29 | SEMA3G | 1.11E-05 | 0.2628125 | 0.0473756 | 0 |
| 30 | FAT2 | 1.49E-07 | 0.195 | 0.0474748 | 0 |
| 31 | METRNL | 8.50E-06 | 0.2484375 | 0.0475732 | 0 |
| 32 | FSTL5 | 8.41E-07 | 0.211875 | 0.0476419 | 0 |
| 33 | SLITRK2 | 1.38E-05 | 0.261875 | 0.0479187 | 0 |
| 34 | GRIA4 | 1.26E-05 | 0.2503125 | 0.0479998 | 0 |
| 35 | KCTD12 | 2.96E-06 | 0.2457813 | 0.0480587 | 0 |
| 36 | QSOX2 | 3.17E-05 | 0.258125 | 0.0480668 | 0 |
| 37 | PREP | 6.19E-05 | 0.26875 | 0.0481573 | 0 |
| 38 | MGAT5 | 4.40E-06 | 0.2453125 | 0.0482849 | 0 |
| 39 | MFGE8 | 5.38E-05 | 0.274375 | 0.048325 | 0 |
| 40 | SLITRK4 | 7.12E-05 | 0.2796875 | 0.0483501 | 0 |
| 41 | GBA | 1.66E-05 | 0.269375 | 0.0484315 | 0 |
| 42 | TIMP4 | 1.55E-05 | 0.2540625 | 0.0485684 | 0 |
| 43 | CHST10 | 7.36E-06 | 0.25125 | 0.0486261 | 0 |
| 44 | BDNF | 4.72E-05 | 0.27625 | 0.048647 | 0 |
| 45 | PDYN | 3.22E-05 | 0.2728125 | 0.048694 | 0 |
| 46 | ADAMTS2 | 3.45E-05 | 0.27 | 0.0487832 | 0 |
| 47 | ST8SIA3 | 3.80E-06 | 0.24625 | 0.0488268 | 0 |
| 48 | APOC3 | 7.47E-05 | 0.28125 | 0.0489095 | 0 |
| 49 | B3GALNT1 | 2.85E-05 | 0.274375 | 0.048914 | 0 |
| 50 | GANAB | 5.38E-05 | 0.27625 | 0.0489428 | 0 |
| Up-regulated proteins |  |  |  |  |  |
| 1 | NEFM | 5.25E-12 | 0.8925 | 0.0320251 | 0 |
| 2 | SERPINA3 | 1.79E-06 | 0.76375 | 0.0437682 | 0 |
| 3 | CHI3L1 | 7.80E-06 | 0.7525 | 0.0438022 | 0 |

**Supplemental Figure S1.**
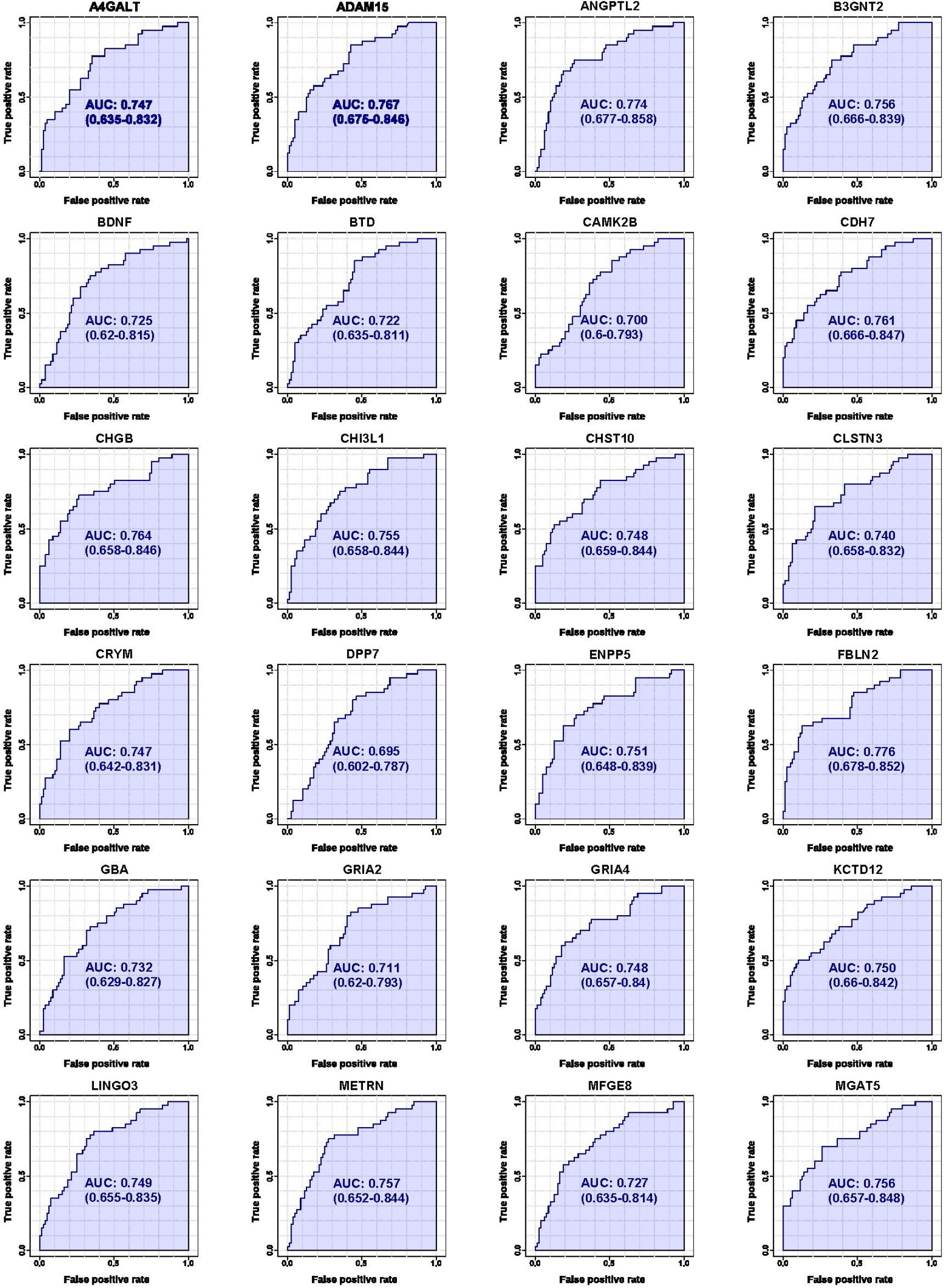
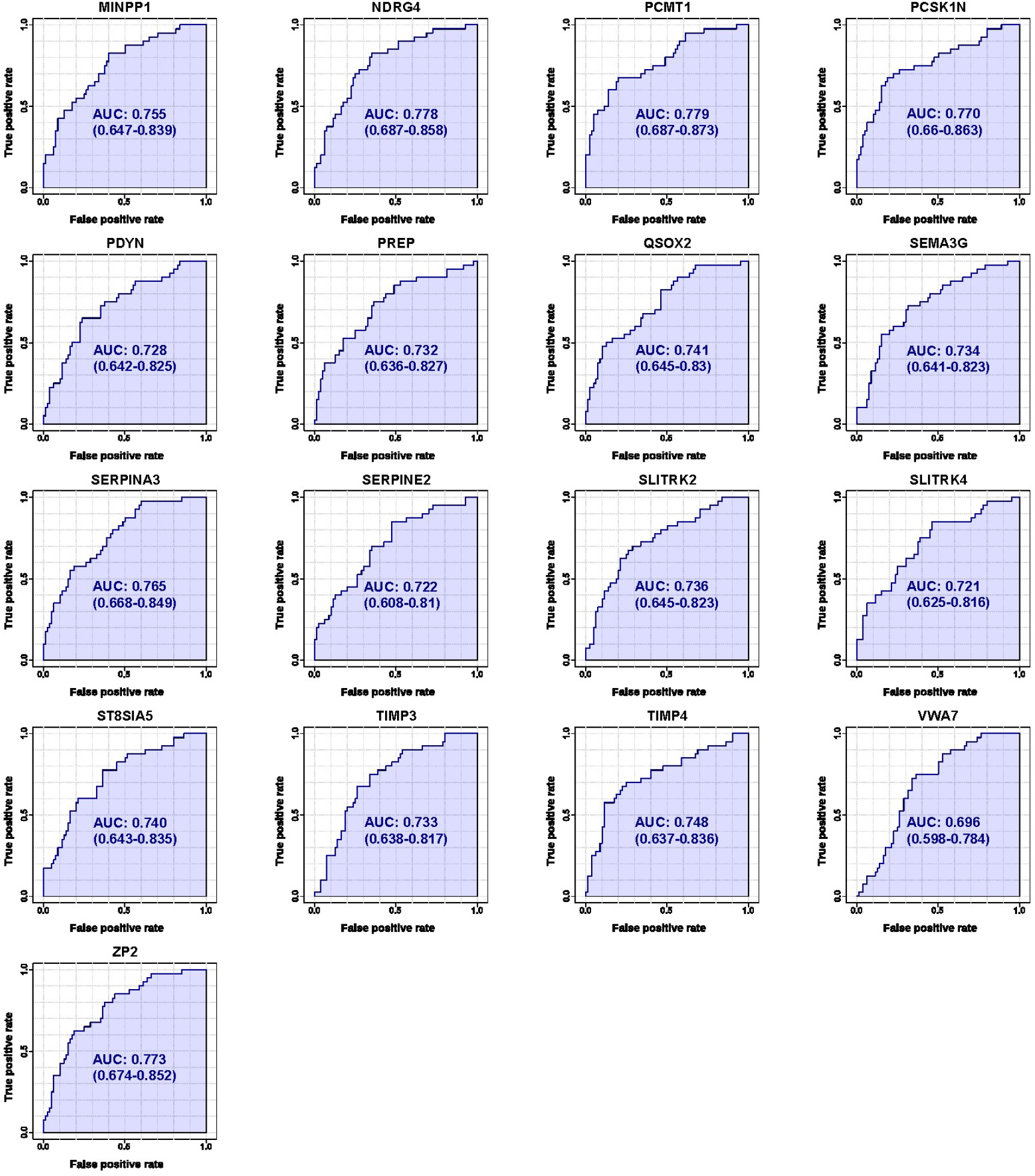
ROC analyses for proteins from the candidate PSP biomarker proteins. The differentially expressed proteins of PSP-specific biomarker candidates were compared with HC plus PD. ROC analysis was generated by bootstrapping. The values in the parenthesis show the lower and upper AUC values of 95% confidence interval. X-axis denotes a false positive rate (1-specificity) and Y-axis denotes a true positive rate (sensitivity).

**Supplemental Figure S2.**
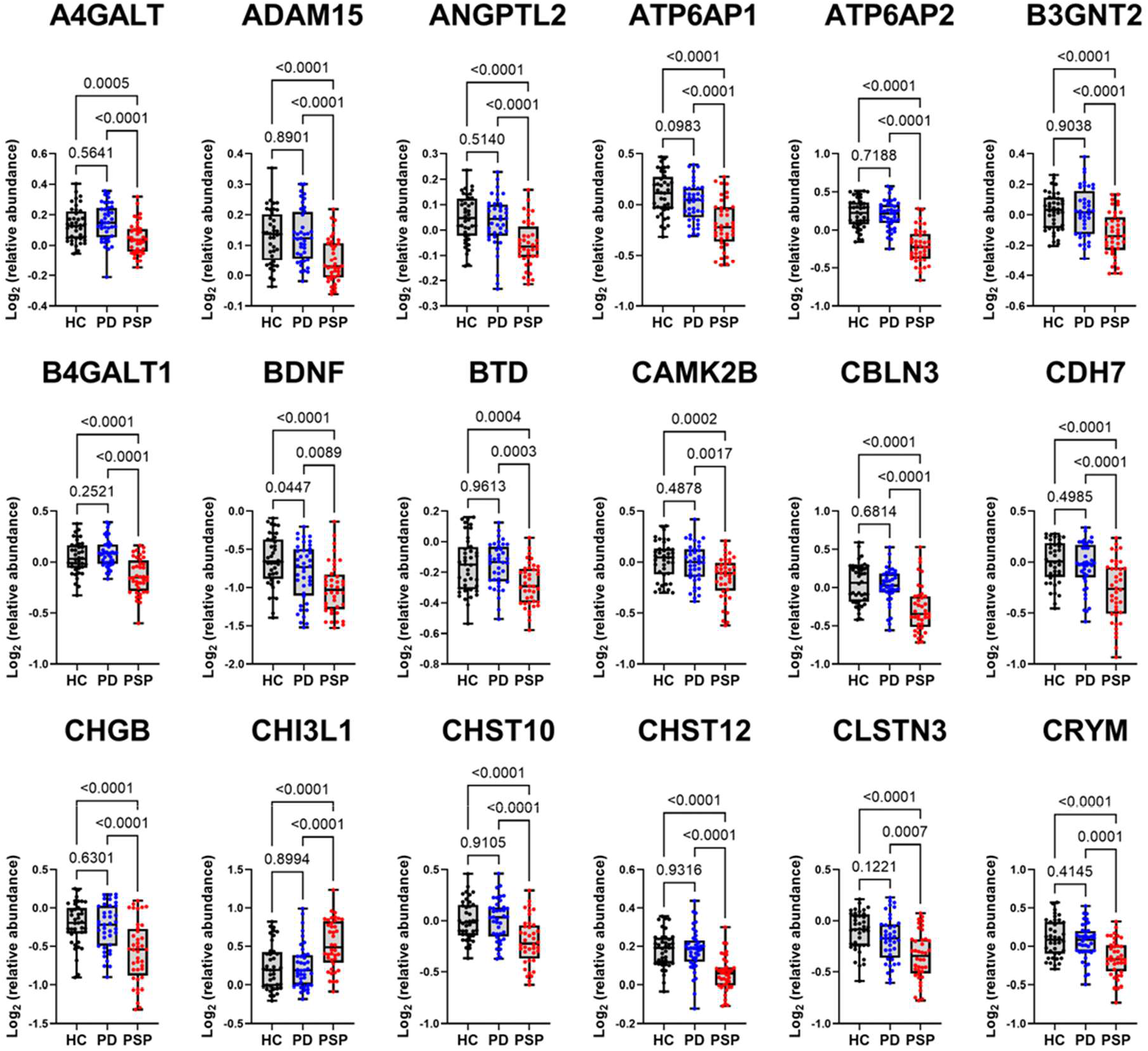
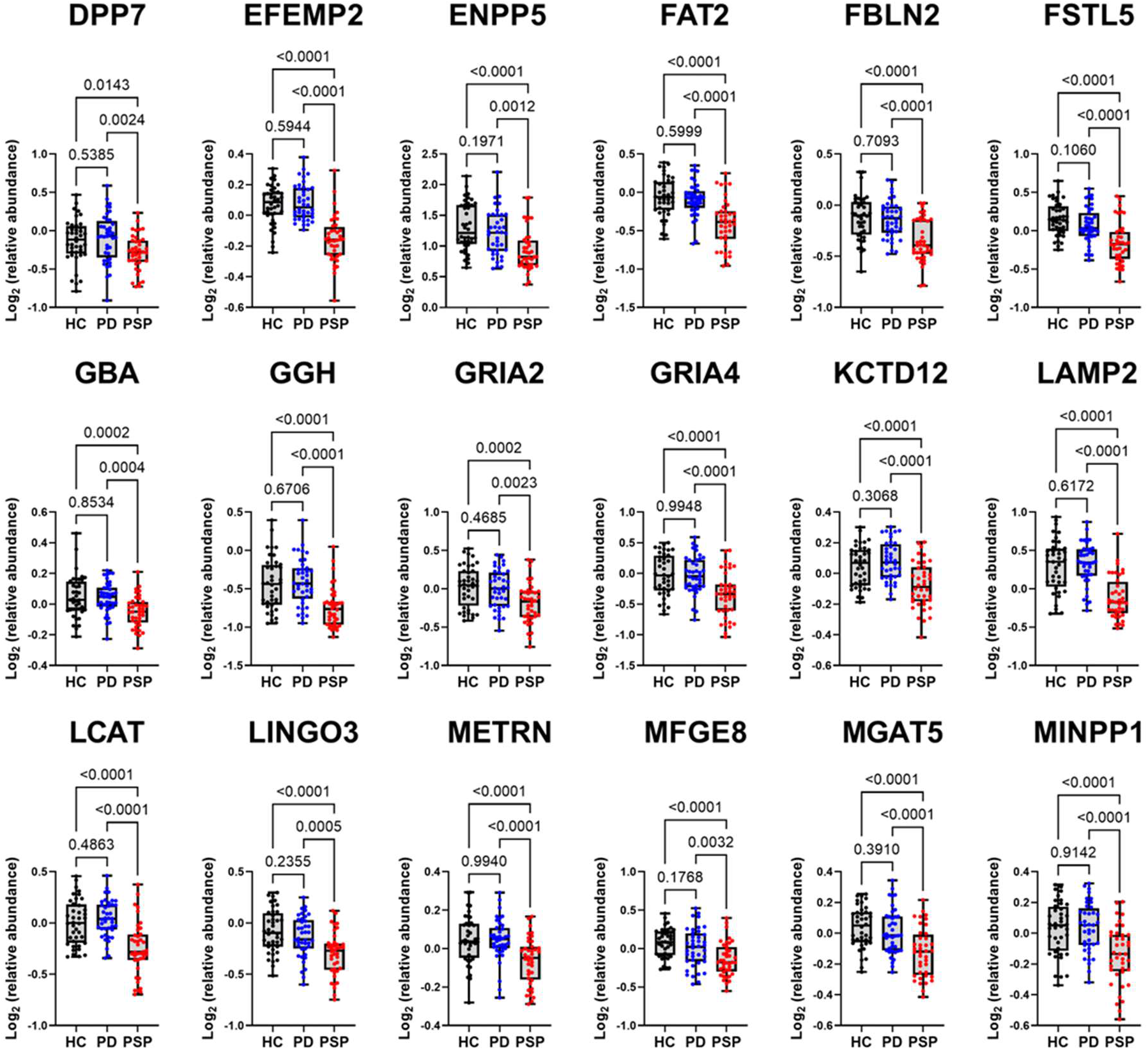
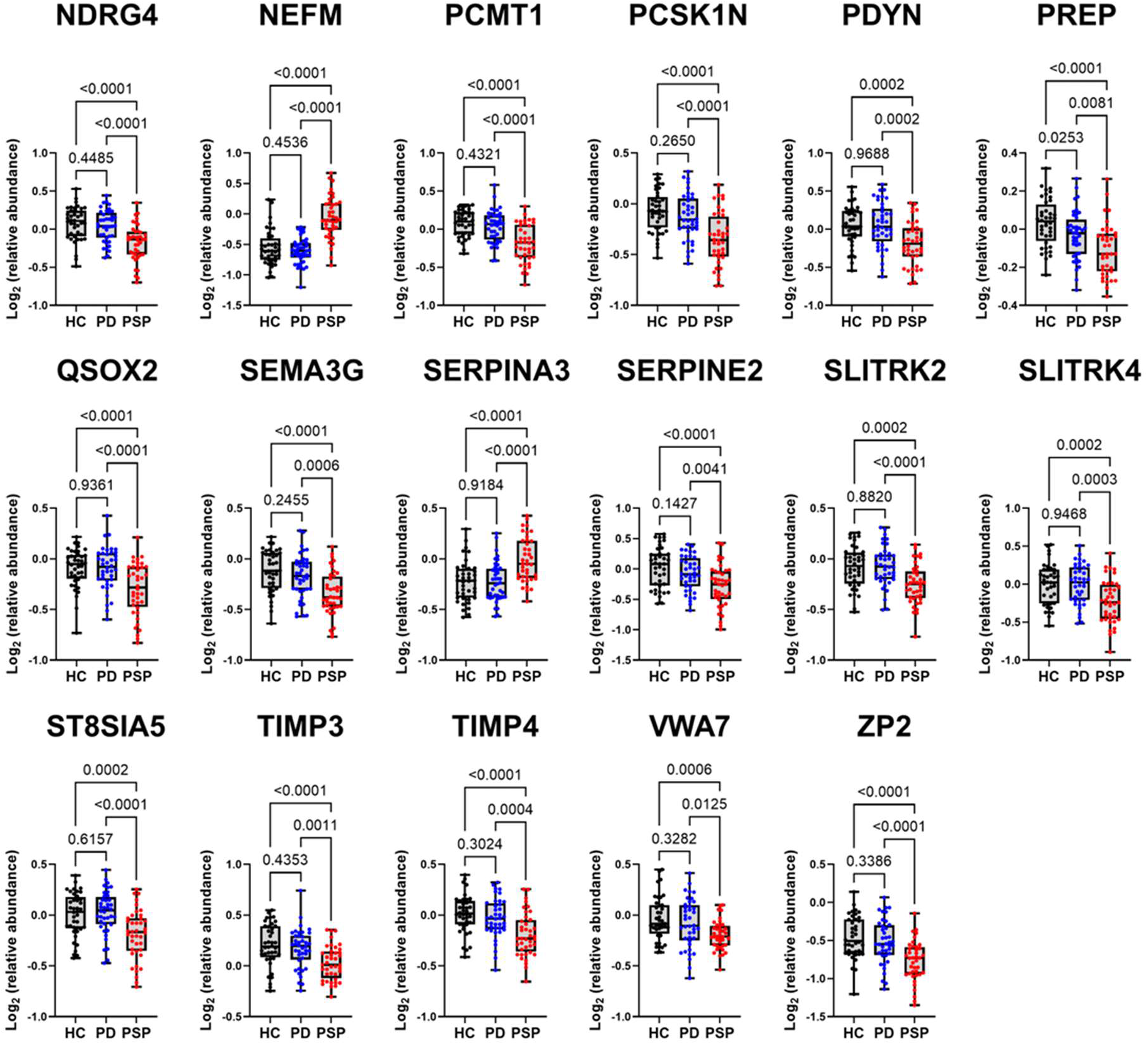
A comparison of the log_2_ relative abundance in PSP and PD plus HC after removing batch effects. The 53 selected PSP-specific biomarker candidates were compared with HC plus PD. Healthy control individuals were shown in black dots, patients with PD were shown in blue dots, and patients with PSP were shown in red dots. The ANOVA test was conducted for statistical analysis between groups (*P*-value shown in figures) using GraphPad Prism 9.

## Notes

### Competing Interest Statement

The authors have declared no competing interest.

